# Tensor decomposition- and principal component analysis-based unsupervised feature extraction to select more reasonable differentially expressed genes: Optimization of standard deviation versus state-of-art methods

**DOI:** 10.1101/2022.02.18.481115

**Authors:** Y-h. Taguchi, Turki Turki

**Affiliations:** Department of Physics, Chuo University, 1-13-27 Kasuga, Bunkyo-ku, 112-8551 Tokyo, JAPAN; Department of Computer Science, King Abdulaziz University, 21589 Jeddah, Saudi Arabia

**Keywords:** tensor decomposition, principal component analysis, feature extraction, standard deviation, differentially expressed genes

## Abstract

**Background:** Tensor decomposition- and principal component analysis-based unsupervised feature extraction were proposed almost 5 and 10 years ago, respectively; although these methods have been successfully applied to a wide range of genome analyses, including drug repositioning, biomarker identification, and disease-causing genes’ identification, some fundamental problems have been identified: the number of genes identified was too small to assume that there were no false negatives, and the histogram of *P*-values derived was not fully coincident with the null hypothesis that principal component and singular value vectors follow the Gaussian distribution.

**Results:** Optimizing the standard deviation such that the histogram of *P*-values is as much as possible coincident with the null hypothesis results in an increase in the number and biological reliability of the selected genes.

**Conclusions:** Tensor decomposition- and principal component analysis-based unsupervised feature extraction are perhaps better than state-of-art methods in regard to predicting differentially expressed genes because they achieve the desired property that the less expressed differentially expressed genes should be less likely selected or even associated with the same amount of logarithmic fold change, although they assume neither negative binomial distribution nor dispersion relation, which is usually assumed in state-of-art methods.

## Background

Identifying differentially expressed genes (DEGs) on the basis of comparative analyses [1, 2] has always been difficult. This challenge is attributable to multiple reasons; however, the primary reason is it being a *large p small n* problem. In a *large p small n* problem, it is difficult to select features based on statistical criteria because a small number of samples (= *n*) have a tendency to lead to low significance; in reality, the obtained *P*-values must be heavily corrected by considering a large number of features (= *p*). This makes it difficult to find features with significance. To resolve this difficulty, many methods specific to gene expression analysis have been proposed. For example, significant analysis microarray (SAM) [3] adds a small amount of constancy to gene expression, thereby avoiding the misidentification of low expressed genes as DEGs. Limma [4] applied a Bayesian strategy to logarithmic gene expression. After high-throughput sequencing (HTS) became popular, *P*-values are attributed to individual genes, assuming that gene expression follows a negative binomial (NB) distribution [5, 6], which is one of the simplest positively valued distributions with a tunable mean and variance. In addition to this, the so-called dispersion relation [5, 6],

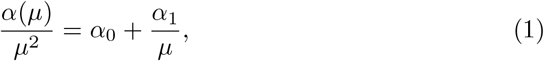

has also been assumed, where *µ* and *α* are the mean and variance, respectively, and *α*_0_ and *α*_1_ are regression coefficients; to our knowledge, eq. (1) is purely empirical and lacks rationalization. Despite these difficulties, many proposed state-of-art methods [5, 6, 7, 8, 9] have been widely employed and used in various stud-ies Contrary to these empirical methods, we proposed tensor decomposition (TD)- and principal component analysis (PCA)-based unsupervised feature extraction (FE) [10] that only assumes that principal component (PC) and singular value vectors (SVVs) obey Gaussian distribution. Despite this simplicity, TD- and PCA-based unsupervised FE have been successfully applied to a wide range of genomic analyses. However, there have been two problems: 1. The histogram of the *P* - values is not fully coincident with the null hypothesis that PC and SVV obey Gaussian distribution and 2. The number of genes selected is too small to have no false negatives. In this paper, we have shown that the optimization of standard deviation (SD) in Gaussian distribution can resolve these problems.

We tried optimizing SD for PCA-based unsupervised FE and applied this to two highly curated data sets––MAQC and SEQC. Then, we tested the optimization of SD for TD-based unsupervised FE and applied it to two more realistic problems: 1. drug repositioning for SARS-CoV-2 and 2. the analysis of gene expression of multiple organs treated with multiple drugs, to which TD-based unsupervised FE without SD optimization was already applied.

## Results

### Outlines of TD and PCA based unsupervised FE

In this section, we have briefly explained the algorithm of PCA- and TD-based unsupervised FE (Fig. 1) before explaining how we could improve them. When a gene expression profile is formatted as a matrix, *x*_*ij*_ ∈ ℝ^*N* × *M*^, which represents the gene expression of the *i*th gene of the *j*th sample, we use PCA-based unsupervised FE. After standardizing *x*_*ij*_ as

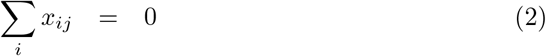

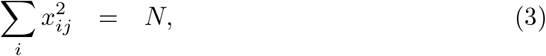

a gram matrix Σ_*j*_ *x*_*ij*_*x*_*i*′*j*_ ∈ ℝ^*N* × *N*^ was diagonalized as

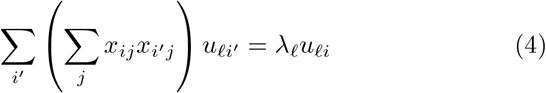

where *u*_*ℓi*_ ∈ ℝ^*N* × *N*^ is the *ℓ*th PC score attributed to gene *i*. The *ℓ*th PC loading attributed to the *j*th sample can be computed as

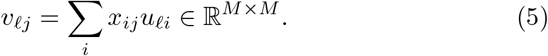

**Figure 1.**
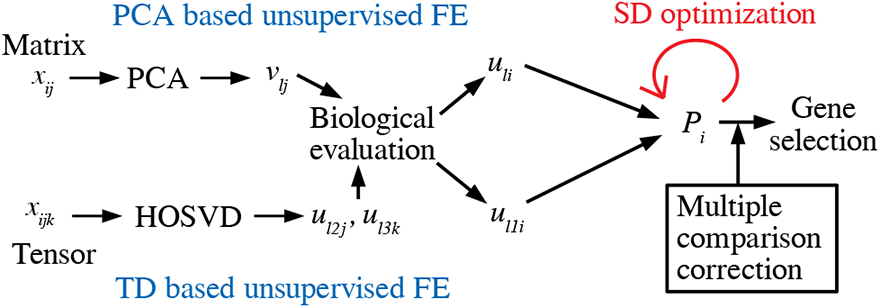
Schematic figure of TD- and PCA-based unsupervised FE with optimized SD.

After identifying *v*_*ℓj*_, which is associated with a desired property, e.g., the district between control and treated samples, we attributed the *P*-values to the gene *i* using the corresponding PC score, *u*_*ℓi*_, as

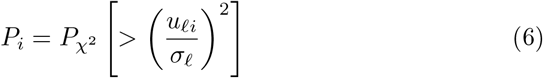

assuming that *u*_*ℓi*_ obeys the Gaussian distribution, where *P*_*χ*2_ [*> x*] is cumulative *χ*^2^ distribution when an argument larger than *x* and *σ*_*ℓ*_ is the SD,

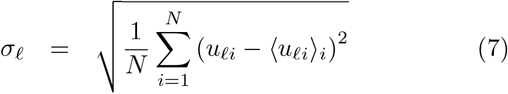

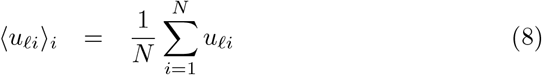

When we have gene expression that is formatted as a tensor, *x*_*ijk*_ ∈ ℝ^*N* × *M* × *K*^, for the expression of the *i*th gene at *j*th sample with the *k*th condition, we used TD-based unsupervised FE. After standardizing *x*_*ijk*_ as

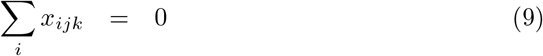

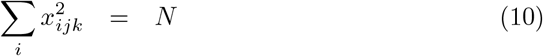

Tucker decomposition of *x*_*ijk*_

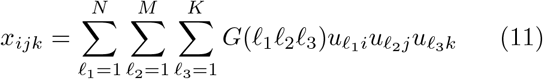

can be computed with a higher order singular value decomposition (HOSVD) [10]. After identifying which 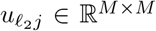 and 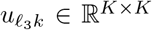 are coincident with the target property, e.g., distinction between control and treated samples specifically under *k*th experimental condition, we try to find *u*_*ℓi*_ ∈ ℝ^*N* × *N*^ associated with *G*(*ℓ*_1_*ℓ*_2_*ℓ*_3_) ∈ ℝ^*N* × *M* × *K*^ having the largest absolute value. Then, the *P*-value is attributed to the *i*th gene as

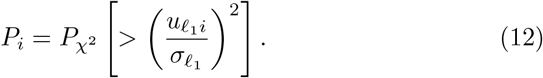

by also assuming that 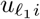 obeys the Gaussian distribution and

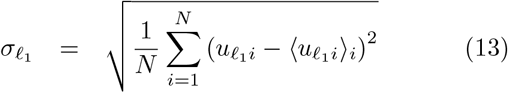

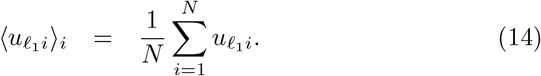

For both PCA- and TD-based unsupervised FE, *P*_*i*_ is corrected with the Benjamini-Hochberg (BH) criterion [10]; further, the *i*th genes associated with adjusted *P*_*i*_ less than the threshold value, which is usually 0.01, are selected.

Although PCA- as well as TD-based unsupervised FE were successfully applied to a wide range of genomic analyses, there were two weak points:

- Too small a number of genes were selected to have no false negatives.
- The histogram of *P*_*i*_ did not fully obey the null assumption that *u*_*ℓi*_ and 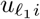 obey the Gaussian distribution.

In this paper, by fixing these two problems, we have tried to establish a new method at least comparable to or even superior to state-of-art methods.

### Trials using highly curated data sets

#### Application to MAQC dataset

Initially, to assess what the problem is, we compared the performance of PCA-based unsupervised FE with DESeq2, a state-of-art method, using the MAQC [11] data set, which has been carefully curated and frequently used for benchmark studies. Figure 2C shows a scatter plot of genes using *u*_1*i*_ and *u*_2*i*_. Figure 2A and B show the PC loading *v*_1*j*_ and *v*_2*j*_; *v*_1*j*_ represents the mean gene expression and *v*_2*j*_ represents the differential expression between universal human reference (UHR) and brain. Occasionally, this reminds us of the horizontal and vertical axes of an MAPlot; the horizontal axis of an MAPlot represents the mean expression of individual genes, typically the mean logarithmic expression,

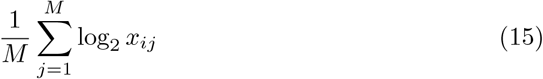

whereas the vertical axis of an MAPlot represents the differential expression between the two classes, typically the mean logarithmic fold change (LFC),

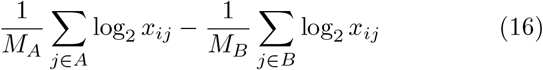

where *M*_*A*_ and *M*_*B*_(= *M* −*M*_*A*_) are sample numbers within one of the two classes, A and B, respectively, and summations are taken within individual classes. As can be seen in Fig. 2D, which represents the contribution of PC loading, *x*_*ij*_ can be expressed almost fully in the 2-dimensional space spanned by the first two PCs. Thus, PCA can derive, in a fully unsupervised manner, something that qualitatively corresponds to an MAPlot (Fig.8), which is usually drawn artificially. In spite of that, unfortunately, the genes selected by the adjusted *P*_*i*_ are too small to have no false negatives (Table 3) and an histogram of *P*_*i*_ is hardly regarded to obey the null hypothesis; the left panel of Fig. 3 shows the histogram of 1−*P*_*i*_, where *P*_*i*_s were computed from *u*_2*i*_ by eq. (6) using *σ*_2_ defined as

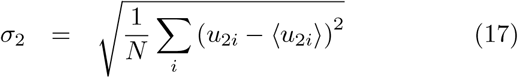

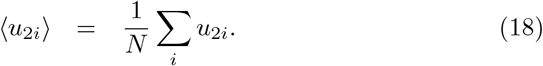

**Figure 2.**
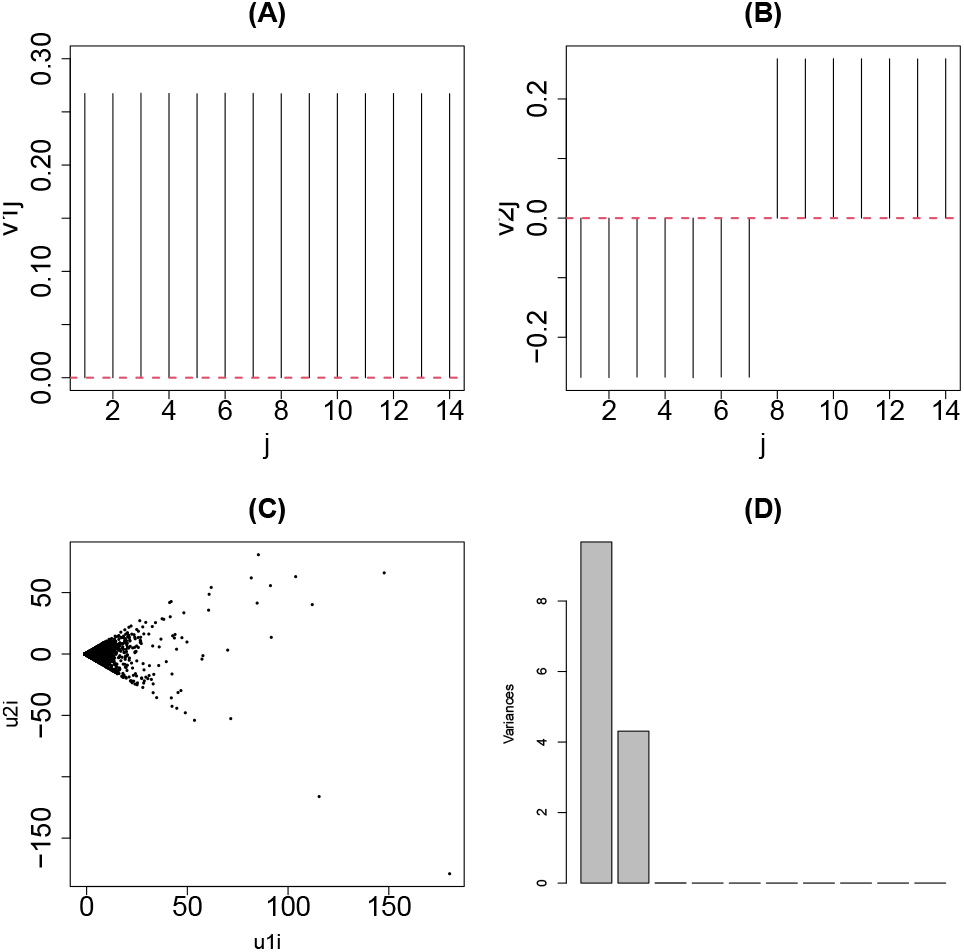
PCA applied to MAQC data. (A) *v*_1*j*_ (B) *v*_2*j*_ (C) Scatter plot of *u*_1*i*_ and *u*_2*i*_ (D) Contributions of individual PCs

**Figure 3.**
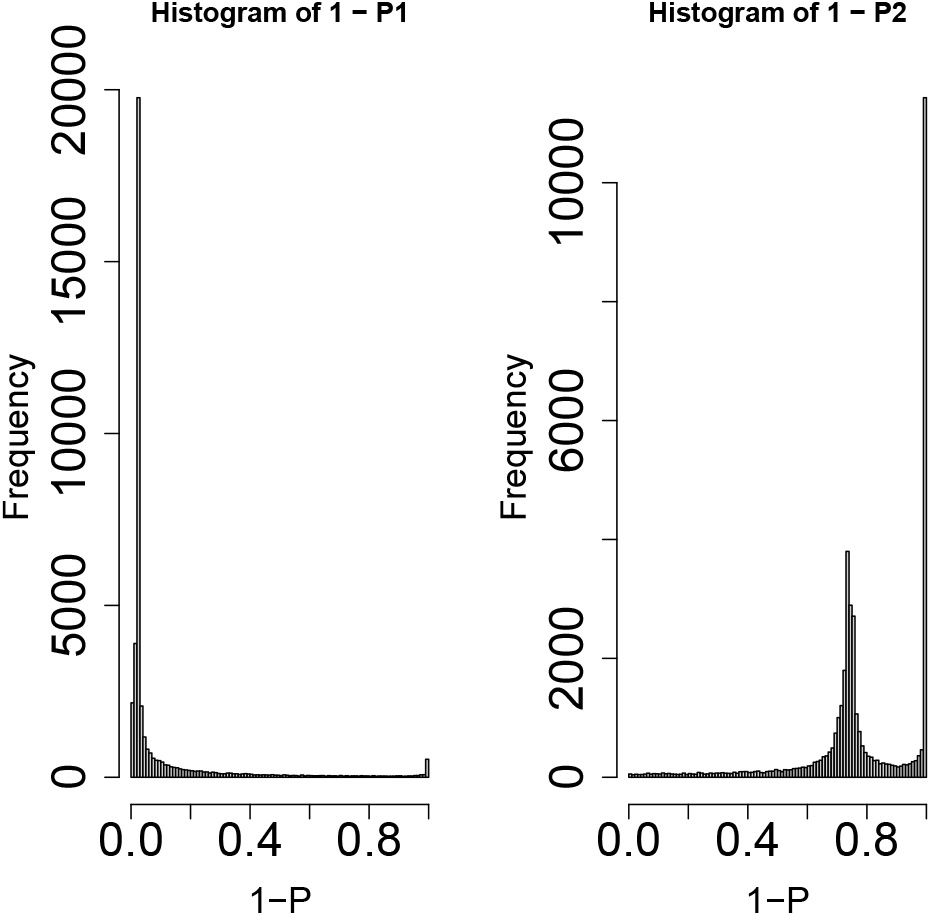
Histogram of 1 − *P*_*i*_ of the MAQC data set with PCA-based unsupervised FE. Left: *P*_*i*_s by eq. (6) using SD *σ*_2_ directly computed from *u*_2*i*_, right: using SD optimized to obey the Gaussian distribution as much as possible.

If 1 − *P*_*i*_ is coincident with the null hypothesis; the histogram of 1 − *P*_*i*_ *<* 1 should have a flat distribution and that of 1 − *P*_*i*_ ∼ 1 should have a sharp peak.

#### Top ranked genes are coincident with DESeq2

To understand the problem of *P*_*i*_s computed by PCA-based unsupervsied FE, we compared *P*_*i*_s computed by PCA-based unsupervised FE with those computed by DESeq2, a state-of-art method. At first, AUC was computed to predict the top 1000 genes based on *P*_*i*_ derived with DESeq2 using *P*_*i*_s computed by PCA-based unsupervised FE; the area under the curve (AUC) was 0.97. Next, in contrast, the AUC was computed to predict the top 1000 genes based on *P*_*i*_ derived with PCA-based unsupervised FE using *P*_*i*_s computed using DE-Seq2; the AUC was 0.98. This indicated that the top-ranked genes were suitably shared between PCA-based unsupervised FE and DESeq2. Thus, the problem of PCA-based unsupervised FE is not the genes’ ranking but the absolute value of *P*_*i*_s.

#### Optimization of SD

Based on the observations at the end of the subsub-section, we arrived at optimizing *σ*_*ℓ*_ such that *u*_*ℓi*_ and *u*_*ℓ*1_ *i* obeyed the Gaussian distribution. Generally, optimizing SD to be fitted to the null hypothesis is not easy. For example, Mudge et al [12] had to assume the equivalence between Type I and II errors, which we cannot assume because of an imbalance of numbers between DEGs and the other genes; typically, DEGs are expected to be minorities. Next, we decided to employ an alternative and more empirical approach. To visualize the idea, we have shown some illustrative examples. Figure 4 shows a historgam of the variable *x*_*i*_ derived from the Gaussian distribution and outliers. If we attribute the *P*-values to the *i*th variable with *x*_*i*_

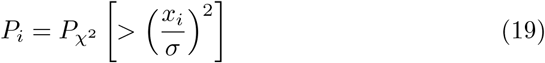

using the SD, *σ*, directly computed by all points

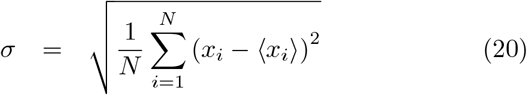

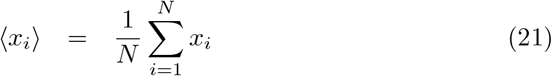

and select outliers associated with adjusted *P* –values *<* 0.01, we cannot select any of the outliers (Table 1); this is because the SD computed, 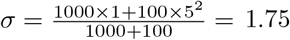, is larger than that of the Gaussian distribution, *σ* = 1, because of outliers. Because *P*_*i*_s computed with *σ* = 1.75 is larger than that with *σ* = 1, it fails to recognize outliers correctly.

**Figure 4.**
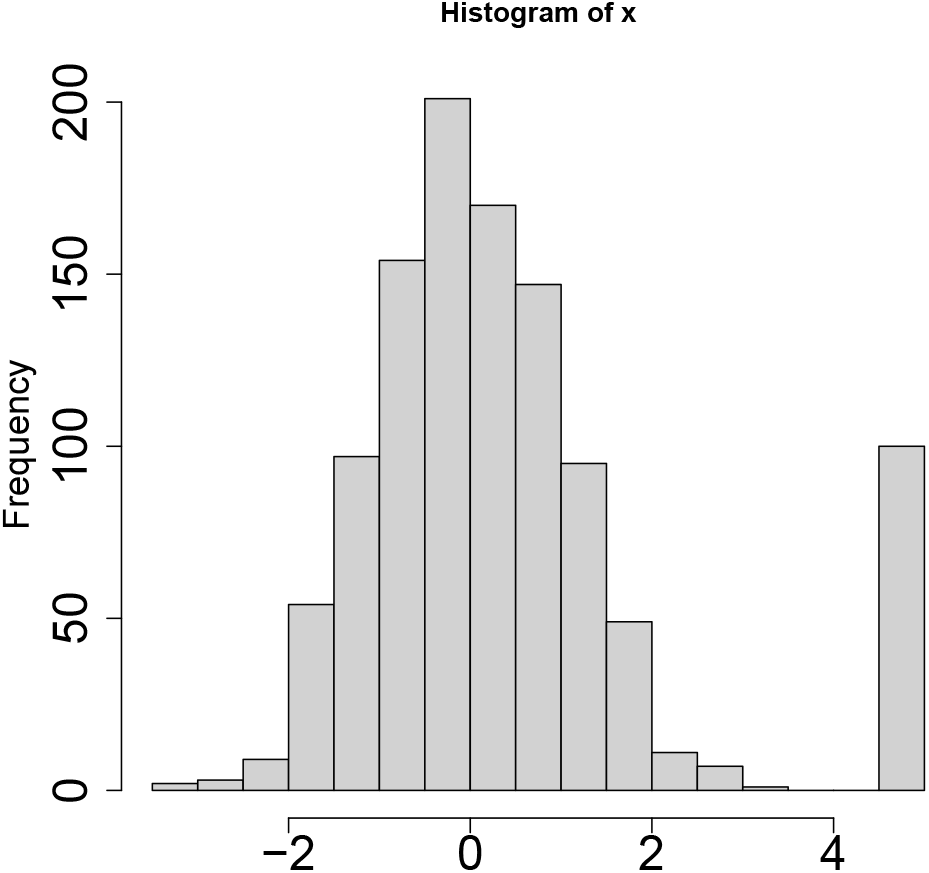
A histogram of Gaussian distribution with outliers.

**Table 1.**
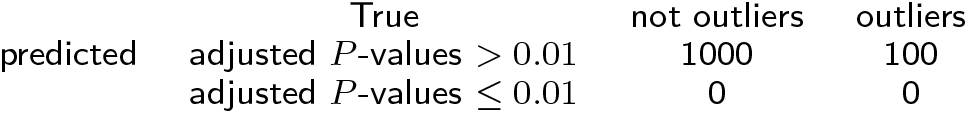
Confusion matrix of the Gaussian distribution with outliers and prediction for *x*_*i*_, the historam for which is given in Fig. 4.

We computed the histogram of 1 − *P*_*i*_, Fig. 5A, which is far being idealized, Fig. 5C, that should have a constant histogram *h*(1 − *P*_*i*_) up to 1 − *P*_*i*_ very close to 1 and has one with a narrow peak near 1 − *P*_*i*_ ∼1. To optimize the SD, we tried to find an optimal SD such that the histogram for those not recognized as outliers was as flat as possible, i.e, obeying the null hypothesis of the Gaussian distribution; we decided to find the optimal SD that results in the most flat *h*(1 − *P*_*i*_) for 1 − adjusted *P*_*i*_ less than threshold value 1 − adjusted *P*_0_ (adjusted *P*_0_ should be small enough). To minimize the SD of binned *h*_*i*_ = *h*(1 − *P*_*i*_), *σ*_*h*_,

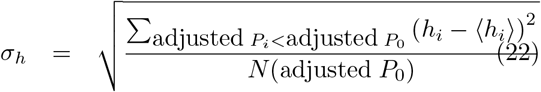

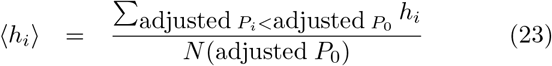

with respect to *σ*, where *N* (adjusted *P*_0_) is the number of *i*s associated with adjusted *P*_*i*_ *>* adjusted *P*_0_, i.e., not recognized as outliers and recognized as a part of the Gaussian distribution. After optimizing *σ*_*ℓ*_, we recomputed *P*_*i*_. Fig. 5A and 5B show the histogram of 1 − *P*_*i*_ using *σ* = 1.75 and optimized SD, respectively; the latter is closer to an idealized histogram of *P*_*i*_, Fig. 5C, than the former.

**Figure 5.**
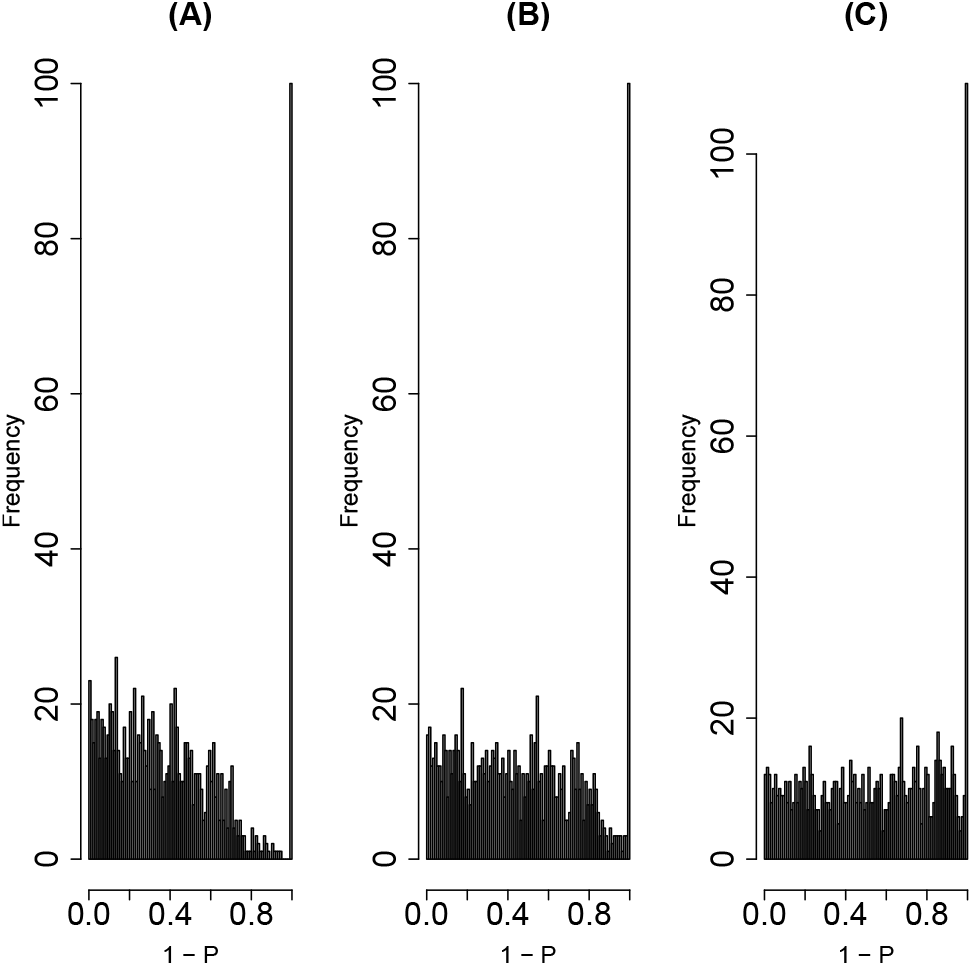
Histograms of 1 − *P*_*i*_, *h*(1 − *P*_*i*_), for (A) *P*_*i*_ computed by eq. (6) with *σ* defined in eq. (20), (B) that with optimized SD, (C) that with true SD, *σ* = 1.

To validate the effectiveness of the optimization of SD, we repeated this procedure 100 times. Figure 6 shows the dependence of *σ*_*h*_ on SD (upper panel) and the comparison between SD in Eq. (20), optimized SD, and SD computed using *i*s for adjusted *P*_*i*_ *<* adjusted *P*_0_ (lower panel). In the lower panel, the optimized SD was approximately 1.2, which is much closer to 1 than 1.75, computed by eq. (20). In addition, the fact that SD computed using *i*s for adjusted *P*_*i*_ *<* adjusted *P*_0_, which is expected to correspond to the Gaussian distribution part in Fig. 4, is almost 1 helps justify our optimization procedure (Fig. 6, lower panel). The reason why SD = 0 with *σ*_*h*_ = 0 in the upper panel of Fig. 6 was not selected as optimal (as having the smallest *σ*_*h*_) is because *σ* = 0 corresponds to nothing selected and is thus meaningless. Using *P*_*i*_ computed by optimized SD, we can discriminate the outliers almost perfectly (Table 2).

**Figure 6.**
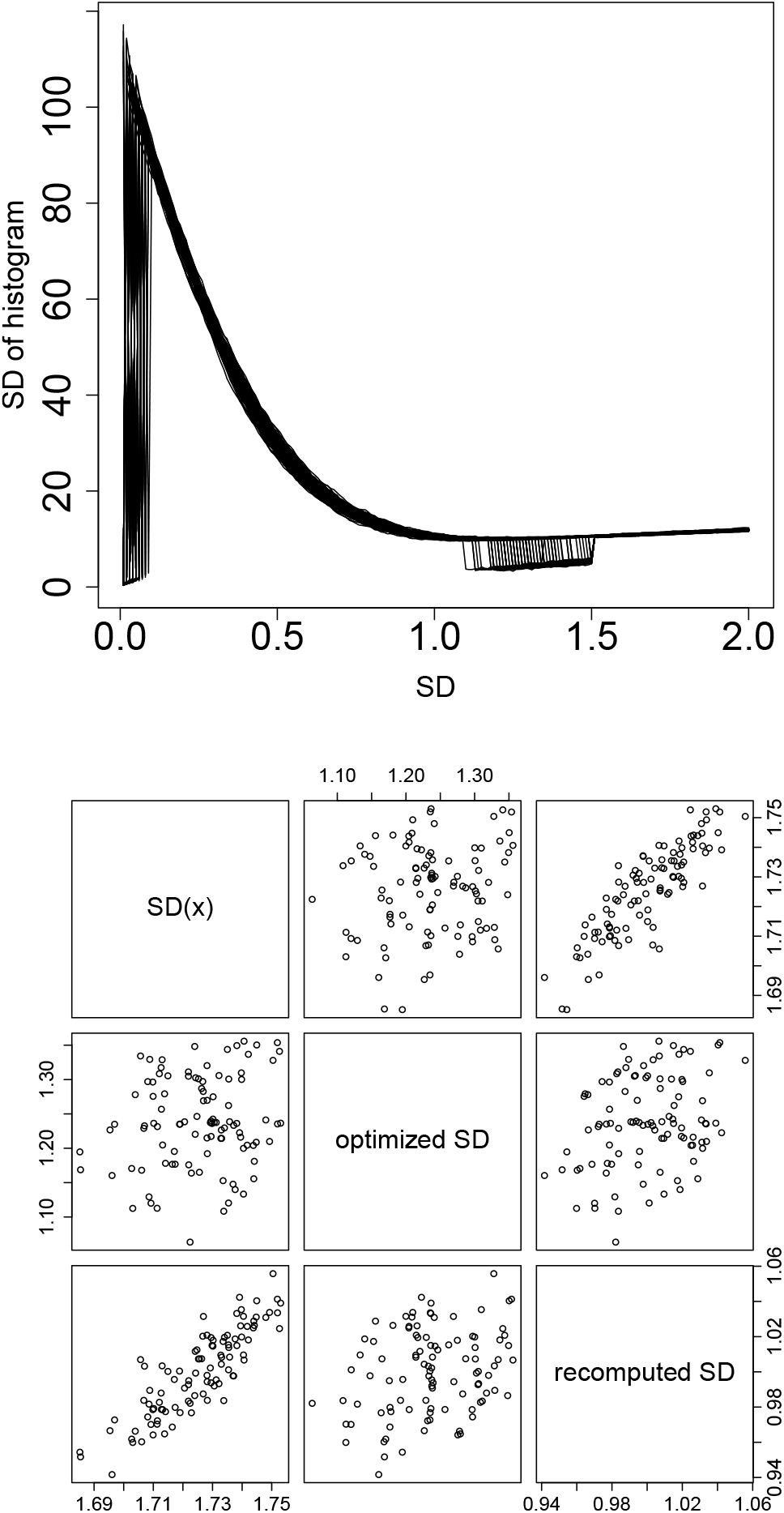
Scatter plot of SDs. Upper: *σ*_*h*_, defined in eq. (22) as a function of SD used for computing *P*_*i*_ in eq. (19). Lower: Scatter plot *σ* of eq. 20, optimized SD, and SD computed using *i*s with adjusted *P*_*i*_ *<* adjusted *P*_0_ (recomputed SD).

**Table 2.**
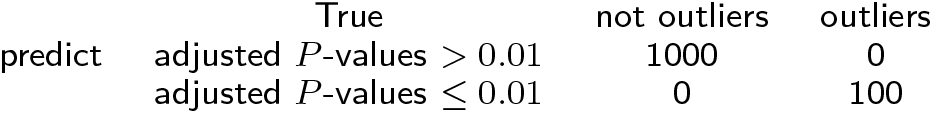
Averaged confusion matrix of Gaussian distribution with outliers and prediction using optimized SD.

|  | True | not outliers | outliers |
| --- | --- | --- | --- |
| predict | adjusted $P$-values $> 0.01$ | 1000 | 0 |
| | adjusted $P$-values $\leq 0.01$ | 0 | 100 |

Next, we applied this strategy to the MAQC data set. Figure 7 shows *σ*_*h*_, defined in eq. (22), as a function of SD to compute *P*_*i*_ in eq. (19) using the MAQC data set; the optimal SD was 0.05557979. It is close to the SD recomputed using *i*s with adjusted *P*_*i*_ *<* adjusted *P*_0_, 0.03871846; moreover, *h*(1 − *P*_*i*_) derived from optimal SD looks more idealized (the right panel of Fig. 3). Thus, the optimal SD improved PCA-based unsupervised FE.

**Figure 7.**
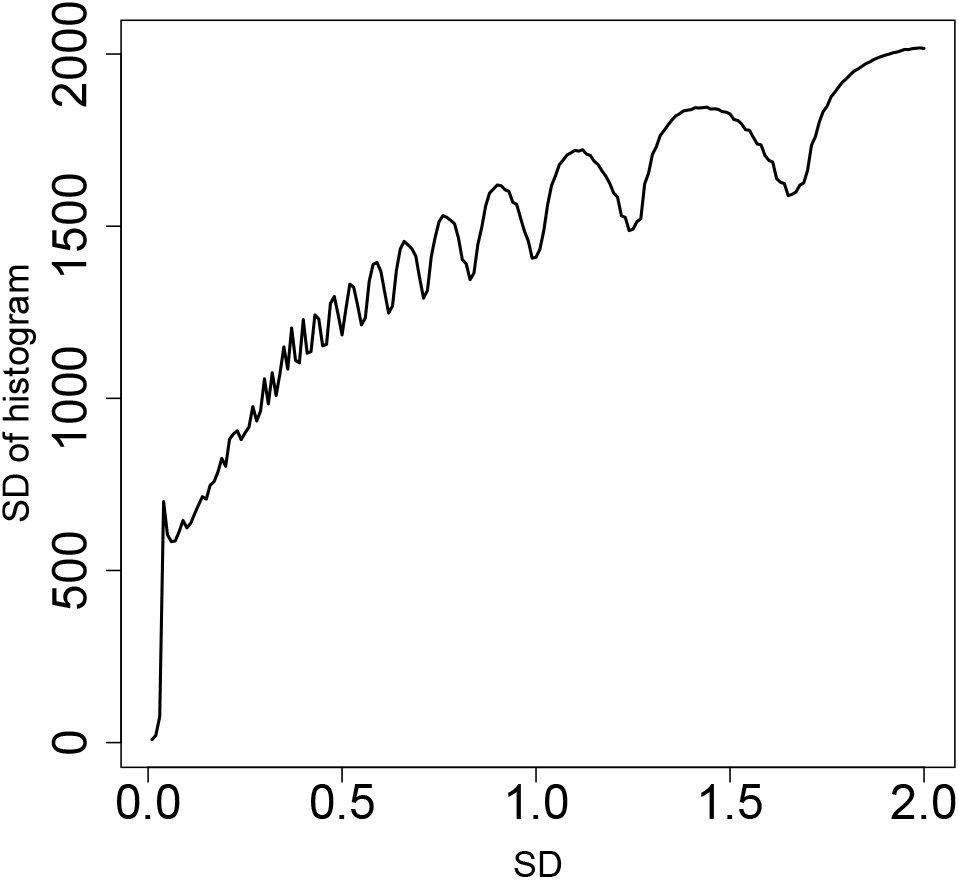
***σ***_*h*_, defined in eq. (22) as a function of SD used for computing *P*_*i*_ in eq. (19) using MAQC data.

Table 3 shows the number of genes selected using DESeq2 (list of genes available as Additional file 1), the original PCA-based unsupervised FE, than by using optimal SD (list of genes available as Additional file 2). Although the number of genes selected by original PCA-based unsupervised FE, 344, is too small to regard no false negatives, that of genes selected by PCA-based unsupervised FE with optimal SD, 12252, is large enough to regard no false negatives. Furthermore, that of DESeq2, 20546, seems to be too large to have no false positives, because it is unlikely true that more than half the genes (40933) are distinctly expressed between the brain and controls.

**Table 3.** The number of genes selected with original PCA-based unsupervised FE, that with optical SD, and DESeq2.

| | adjusted $P_i$ | |
| --- | --- | --- |
| | $> 0.01$ | $\leq 0.01$ |
| PCA based unsupervised FE |  |  |
| original ( without optimal SD) | 40589 | 344 |
| with optimal SD | 28681 | 12252 |
| DESeq2 | 8789 | 20546 |

#### Less expressed genes are less likely to be DEGs

Figure 8 shows the selected genes in MAPlot. Although we assumed neither NB distribution nor dispersion relation, eq. (1), the distribution of selected genes in the MAPlot is reasonable; genes with the same LFC (vertical axis) are less likely selected when associated with smaller mean expression (horizontal axis). Although this property is explicitly assumed in DESeq2 with dispersion relation, eq. (1), PCA-based unsupervised FE seems to possess the property without assuming dispersion relation explicitly (see the Discussion section). On the other hand, DESeq2 selects too many genes and is less likely reasonable. This suggests that PCA-based unsupervised FE with optimized *σ*_*ℓ*_ is a promising method.

**Figure 8.**
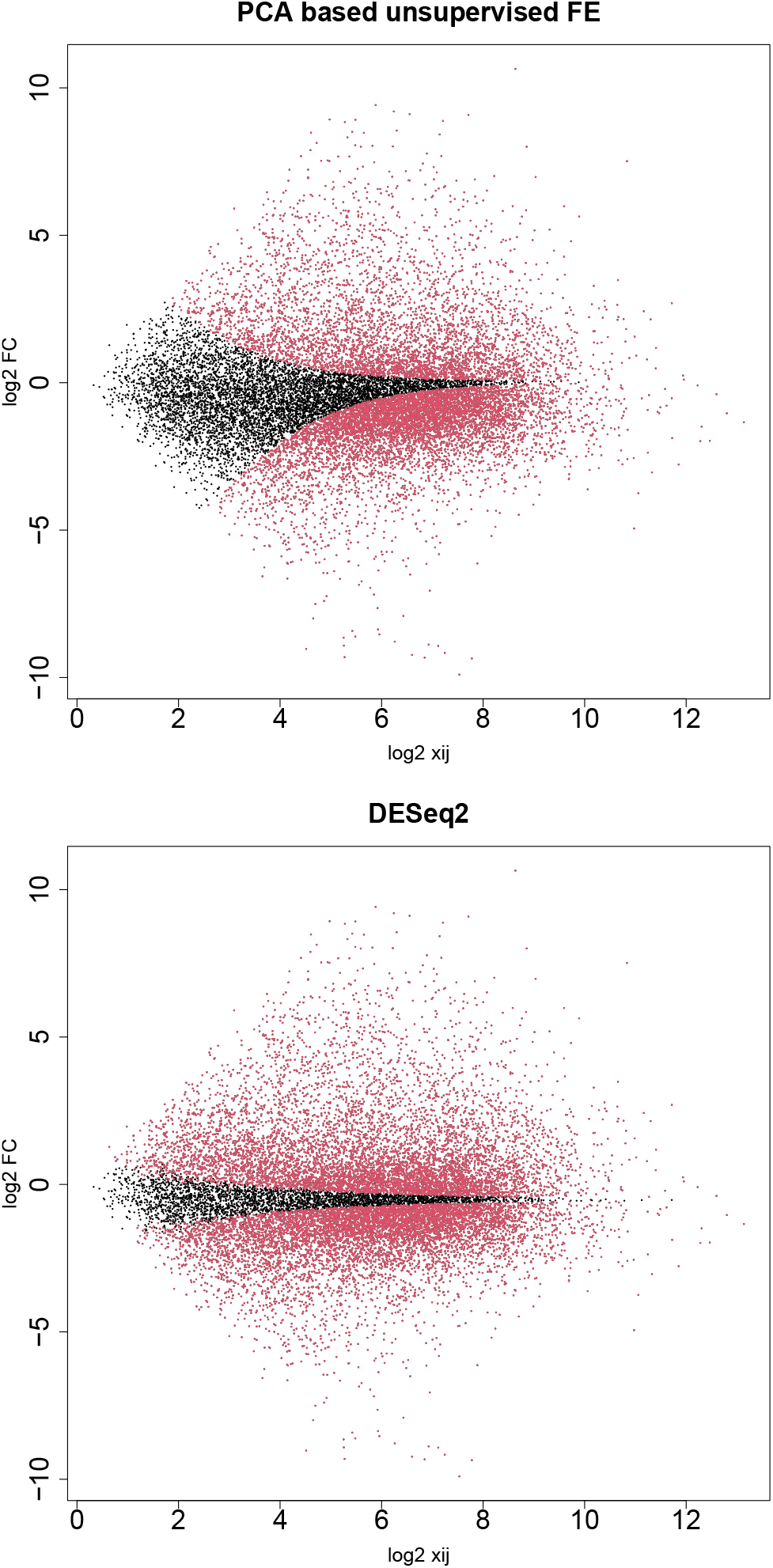
MAPlot with selected genes colored in red. Upper: PCA-based unsupervised FE with optimized SD, lower: DESeq2.

#### Confirmation using the SEQC dataset

To see if it occurs only occasionally, we repeated all computations on as many as 13 data sets in SEQC [13], which is yet another curated data set. Coincidence between DESeq2 and PCA-based unsupervised FE (Fig. 9), a reasonable number of selected genes (∼10^3^, Fig. 10), and a lower opportunity of less expressed genes to be DEGs (Fig. 11) are also observed, as in the case of MAQC. In addition to this, although the number of genes selected by DESeq2 are too large (∼ 10^4^) and heavily dependent upon sample numbers (∼10^3^ for the smallest sample number ∼10^0^), that by PCA-based unsupervised FE is not and is always ∼ 10^3^, regardless of sample numbers. Thus, PCA-based unsupervised FE is seemingly superior to DESeq2.

**Figure 9.**
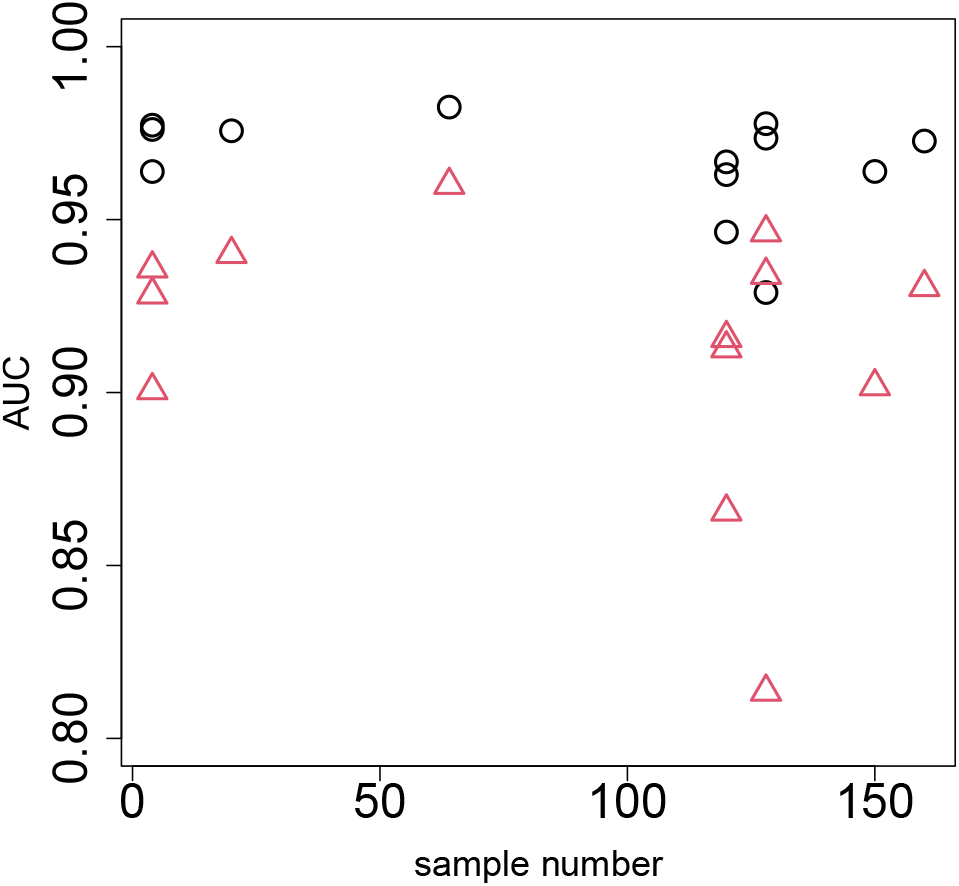
Coincidence of top-ranked genes between DESEq2 and PCA-based unsupervised FE using the SEQC data set. Open circles: AUC when *P*-values computed by PCA-based unsupervised FE with optimized SD discriminates top 1000 genes ranked by *P*-values computed by DESeq2. Open red triangles: AUC when *P*-values computed by DESeq2 discriminating top 1000 genes ranked by *P*-values computed by PCA-based unsupervised FE with optimized SD.

**Figure 10.**
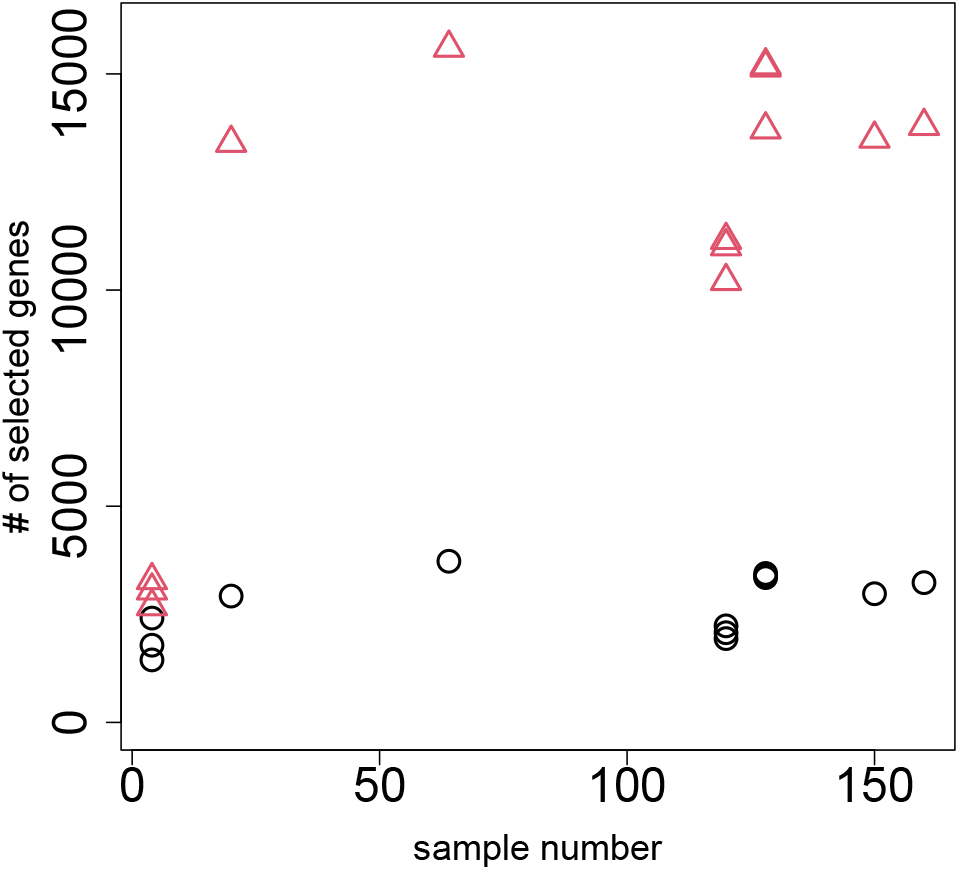
Dependence of the number of DEGs on sample numbers using the SEQC data set. Open circles: the number of genes selected by PCA-based unsupervised FE with optimized SD. Open red triangles:the number of genes selected by DESeq2.

**Figure 11.**
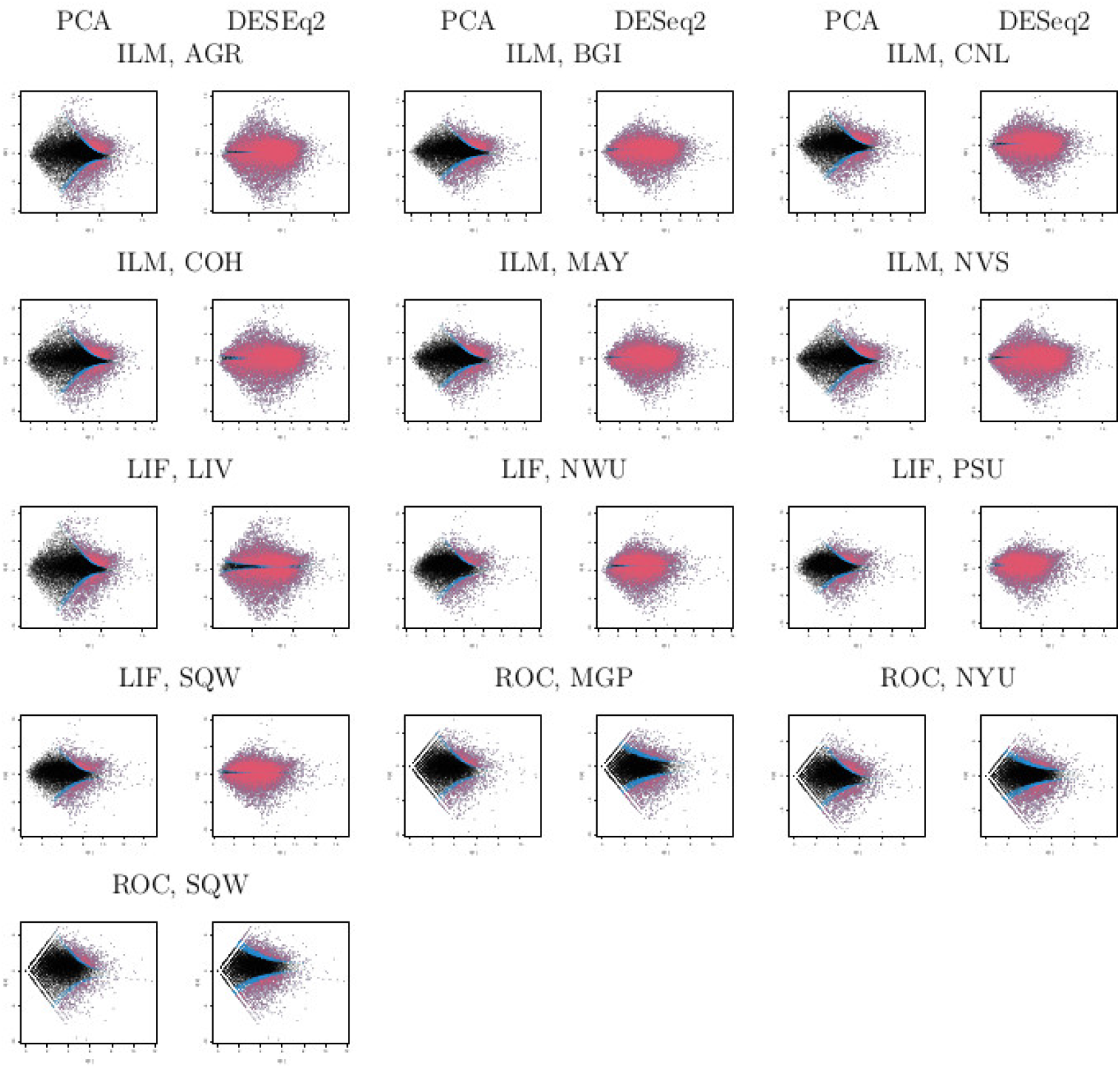
MAPlot for SEQC. PCA-based unsupervised FE with optimized SD: the first, third, and fifth columns, DESeq2: the second, forth, and sixth columns. Three character IDs represent platform and sites. Blue: genes associated with adjusted *P*-values less than 0.1 but greater than 0.01. Red: genes associated with adjusted *P*-values less than 0.01.

#### Biological validation

Based on the above results, PCA-based unsupervised FE is seemingly better than DESeq2. Nonetheless, PCA-based unsupervised FE can select a reasonable number of genes regardless of sample numbers (Fig. 10), and less expressed genes are unlikely to be DEGs when genes are selected by PCA-based unsupervised FE with optimized SD (Figs. 8 and 11), even without assuming NB distribution and dispersion relations, eq. (1), which DESeq2 requires, if the selected genes are not biological, it is meaningless. To evaluate the selected genes biologically, we uploaded the genes selected using MAQC to Enrichr. As can be seen in Fig. 12, the genes selected by PCA-based unsupervised FE were better than those selected by DESeq2 (Full list of enrichment analysis is available in Additional files 1 and 2).

**Figure 12.**
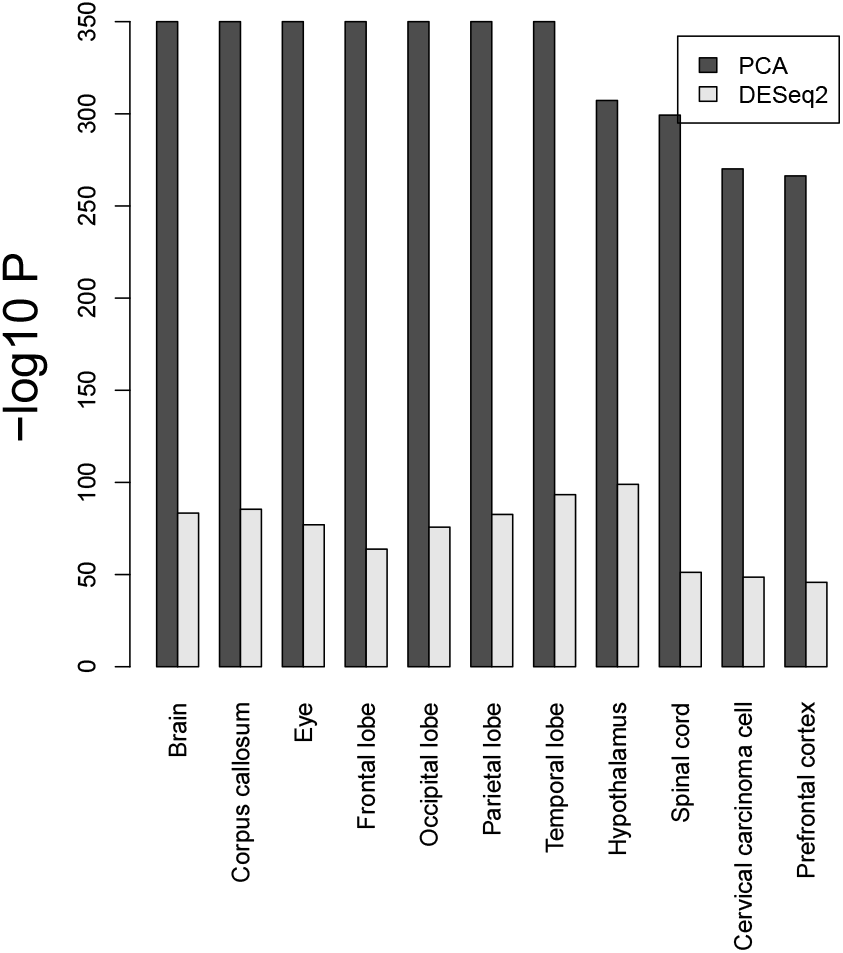
Enrichment analysis of the selected genes, whose numbers in Table 3. *P*-values are adjusted *P*-values (based upon “Jensen Tissues” category in Enrichr). Seven terms associated with − log_10_ *P* = 350 are linked with ℞, since *P* = 0.

One may still wonder the other state-of-art methods might be better than PCA-based unsupervised FE. To deny this possibility, we biologically evaluated the genes selected for MAQC using edgeR [6] (full list of enrichment analysis available in Additional file 3), voom [8] (full list of enrichment analysis available in Additional file 4), and NOISeq [9] (full list of enrichment analysis available in Additional file 5); it is obvious that these three methods are even inferior to DESeq2 biologically (Fig. 13).

**Figure 13.**
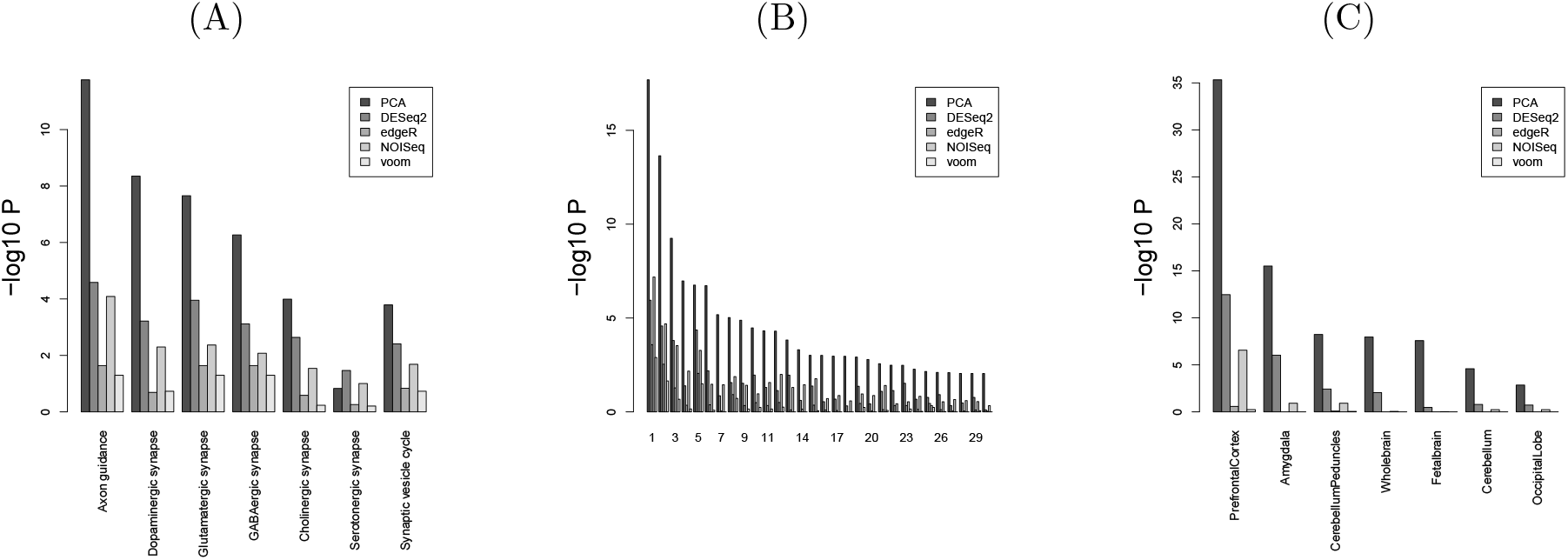
Enrichment analysis for MAQC with other methods in Enrichr. (A) KEGG (B) GO BP (C) Human gene atlas. Numbers in (B) correspond to 1. “axonogenesis,” 2. “axon guidance,” 3. “axon development,” 4.”regulation of axonogenesis,” 5. “synapse organization,” 6. “modulation of chemical synaptic transmission,” 7. “positive regulation of axonogenesis,” 8.”modulation of excitatory postsynaptic potential,” 9. “regulation of axon extension,” 10. “positive regulation of synaptic transmission,” 11. “axon extension,” 12. “negative regulation of axonogenesis,” 13. “chemical synaptic transmission,” 14. “signal release from synapse,” 15. “synapse assembly,” 16. “regulation of neuronal synaptic plasticity,” 17. “positive regulation of axonextension,” 18. “regulation of trans-synaptic signaling,” 19. “positive regulation of excitatory postsynaptic potential,” 20. “negative regulation of axon extension,” “regulation of synapse assembly,” 22.”retrograde axonal transport,” 23. “synaptic vesicle endocytosis,” 24.”synaptic transmission, GABAergic,” 25. “synaptic transmission, glutamatergic,” 26.”regulation of long-term synaptic potentiation,” 27. “regulation of axon extension involved in axon guidance,” 28. “synaptic membrane adhesion,” 29. “regulation of synaptic transmission, glutamatergic,” 30. “regulation of postsynaptic neurotransmitter receptor activity.” *P*-values are adjusted *P*-values.

### Drug discovery for SARS-CoV-2

Although we have demonstrated that PCA-based unsupervised FE with optimized SD can outperform other state-of-art methods in highly curated data, one might wonder that it is not the case for a realistic and more noisy case. To check if PCA-based unsupervised FE with optimized SD can outperform DESeq2 in more realistic data sets, we considered the drug repositioning of SARS-CoV-2, to which we applied TD-based unsupervised FE [14] and its kernelized version [15].

In our implementation, we employed HOSVD to obtain the tensor decomposition, eq. (11); because HOSVD is equivalent to SVD applied to a matrix obtained by unfolding a tensor, we can obtain the identical *u*_*ℓi*_ independent of which of PCA or HOSVD is used; SD used in eq. (12) can be optimized too. Next, we applied the optimization of SD and could select 3627 genes associated with adjusted *P*-values of less than 0.1 (list of genes available as Additional file 6), which is a much higher number of genes than 163 genes than that selected in previous studies [14, 15].

#### Overlap with human genes known to interact with SARS-CoV-2 protein

We evaluated the selected 3627 genes based on the overlap with the human genes known to interact with SARS-CoV-2, as has been done in previous studies [14, 15] (Fig. 14). It is obvious that TD-based unsupervised FE with an optimized SD can outperform kernel TD-based unsupervised FE, original (without optimized SD) TD-based unsupervised FE as well as DESeq2 (list of overlap available in Additional File 7). Thus, it is indeed an outstanding method.

**Figure 14.**
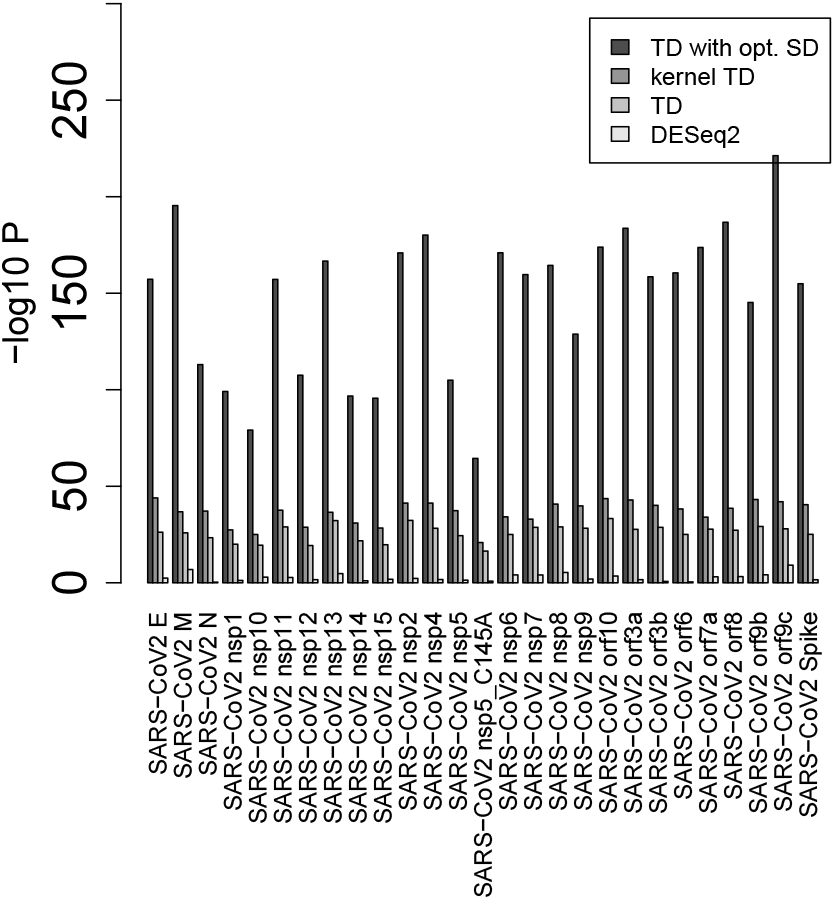
***P*-values computed by Fishers’ exact test to evaluate the overlap between human genes known to interact with SARS-Cov-2 proteins** and genes selected by various methods. DESeq2 is only for A549 cell lines.

#### Drug repositioning

We also tried drug discovery using the genes selected by TD-based unsupervised FE with optimized SD. See Table 4 (Full list of drug repositioning available as Additional file 6). The first one, imatinib, was once identified as a promising drug toward COVID-19, although it was rejected later [16]. The second one, apratoxin A, was reported to be a promising compound based on its protein binding affinity [17]. The third and fourth one, doxycycline, was supposed to be a promising drug to-ward COVID-19 [18]. The seventh one, trovafloxacin, was reported to be a promising compound based on its protein binding affinity [19]. The eighth one, doxorubicin, was also reported to be a promising compound based on its protein binding affinity [20]. The ninth one, cisplatin, and the tenth one, carboplatin, were proposed as a result of drug repositioning [21]. Seven of the nine compounds identified as the top 10 compounds have been previously reported as drugs toward SARS-CoV-2.

**Table 4.** Drug perturbations from GEO down

| Rank | Term | Overlap | P-value | Adjusted P-value | Odds Ratio |
| --- | --- | --- | --- | --- | --- |
| 1 | imatinib (glivec) 123596 human GSE12211 sample 2518 | 316/442 | 7.81E-137 | 7.06E-134 | 12.3 |
| 2 | apratoxin A 6326668 human GSE2742 sample 3071 | 279/389 | 3.77E-121 | 1.57E-118 | 12.3 |
| 3 | doxycycline DB00254 human GSE2624 sample 3074 | 294/425 | 5.22E-121 | 1.57E-118 | 10.9 |
| 4 | doxycycline DB00254 human GSE2624 sample 3077 | 278/391 | 3.83E-119 | 8.64E-117 | 11.9 |
| 5 | grepafloxacin 72474 human GSE9166 sample 2627 | 320/495 | 5.62E-119 | 1.02E-116 | 8.96 |
| 6 | clinafloxacin 60063 human GSE9166 sample 2625 | 309/470 | 8.04E-118 | 1.21E-115 | 9.38 |
| 7 | trovafloxacin 62959 human GSE9166 sample 2629 | 302/459 | 3.05E-115 | 3.94E-113 | 9.38 |
| 8 | doxorubicin, 2xEC50, 5 d 31703 human GSE6930 sample 3265 | 314/493 | 4.76E-114 | 5.37E-112 | 8.57 |
| 9 | cisplatin DB00515 human GSE6410 sample 2532 | 239/315 | 1.06E-112 | 1.07E-110 | 15.1 |
| 10 | carboplatin DB00958 human GSE7035 sample 3060 | 284/422 | 4.57E-112 | 4.13E-110 | 9.99 |

See Table 5. The first, fourth, and tenth one, estradiol, was reported as a promising compound [22]. The second one, tamoxifen, was reported to inhibit SARS-CoV-2 infection by suppressing viral entry [23]. The third one, apratoxin A, has been listed in Table 4, too. The fifth one, MK-886, was reported to be an inhibitor of 3CL protease [24], although its efficiency was limited to 40 %. The sixth one, IFN-alphacon1, was reported to be an inhibitor of SARS-CoV [25] but not for SARS-CoV-2. The seventh one, arachidonic acid, was generally expected to inhibit SARS-CoV-2 infection [26]. The eighth one, arsenic, was also generally expected to act against the RdRp of coronavirus [27]. The ninth one, metoprolo, was reported to be a promising drug toward COVID-19 [28]. Thus, all the top 10 compounds were reported to be promising.

**Table 5.** Drug perturbations from GEO up

| Rank | Term | Overlap | P-value | Adjusted P-value | Odds Ratio |
| --- | --- | --- | --- | --- | --- |
| 1 | estradiol 5757 human GSE4668 sample 3063 | 276/367 | 1.26E-128 | 1.14E-125 | 14.74 |
| 2 | tamoxifen DB00675 human GSE4025 sample 2820 | 271/361 | 6.30E-126 | 2.85E-123 | 14.61 |
| 3 | apratoxin A 6326668 human GSE2742 sample 3068 | 278/389 | 4.61E-120 | 1.12E-117 | 12.16 |
| 4 | estradiol DB00783 human GSE4668 sample 2727 | 261/350 | 4.96E-120 | 1.12E-117 | 14.19 |
| 5 | MK-886 CID 3651377 human GSE3202 sample 3193 | 268/368 | 5.29E-119 | 9.59E-117 | 12.98 |
| 6 | IFN-alphacon1 DB05258 human GSE5542 sample 2474 | 242/313 | 2.21E-117 | 3.34E-115 | 16.41 |
| 7 | Arachidonic acid DB04557 human GSE3737 sample 3171 | 277/395 | 2.80E-116 | 3.63E-114 | 11.39 |
| 8 | ARSENIC 5359596 human GSE6907 sample 3529 | 276/394 | 1.15E-115 | 1.30E-113 | 11.35 |
| 9 | metoprolol DB00264 human GSE3356 sample 2786 | 306/469 | 2.67E-115 | 2.68E-113 | 9.16 |
| 10 | estradiol 5757 human GSE4668 sample 3062 | 245/325 | 1.92E-114 | 1.74E-112 | 14.75 |

On the other hand, for DESeq2, see Table 6 (full list of drug repositioning is available in Additional file 8), The use of the second and third one, dexamethasone, resulted in lower 28-day mortality among those who received either invasive mechanical ventilation or oxygen alone at randomization but not among those receiving no respiratory support. [29], The seventh one, metformin, suppressed SARS-CoV-2 in cell culture [30]. The eighth one, etanercept, significantly decreased the risk of developing COVID-19 in patients with rheumatoid arthritis or spondyloarthropathies [31]. The tenth one, lipopolysaccharide, is not a compound but a bacterial protein reported to bind to the SARS-CoV-2 spike protein [32].

**Table 6.**
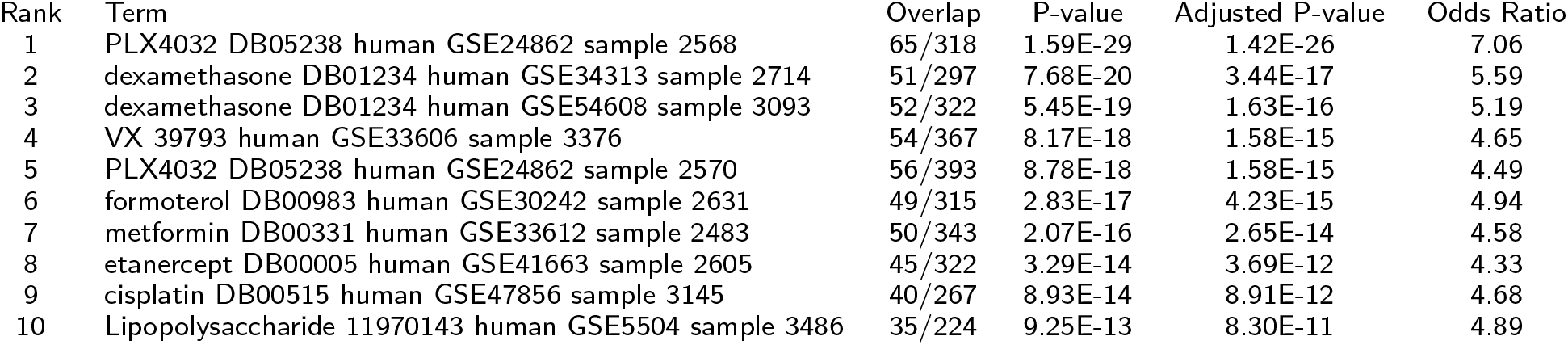
Drug perturbations from GEO down for A549 by DESeq2

| Rank | Term | Overlap | P-value | Adjusted P-value | Odds Ratio |
| --- | --- | --- | --- | --- | --- |
| 1 | PLX4032 DB05238 human GSE24862 sample 2568 | 65/318 | 1.59E-29 | 1.42E-26 | 7.06 |
| 2 | dexamethasone DB01234 human GSE34313 sample 2714 | 51/297 | 7.68E-20 | 3.44E-17 | 5.59 |
| 3 | dexamethasone DB01234 human GSE54608 sample 3093 | 52/322 | 5.45E-19 | 1.63E-16 | 5.19 |
| 4 | VX 39793 human GSE33606 sample 3376 | 54/367 | 8.17E-18 | 1.58E-15 | 4.65 |
| 5 | PLX4032 DB05238 human GSE24862 sample 2570 | 56/393 | 8.78E-18 | 1.58E-15 | 4.49 |
| 6 | formoterol DB00983 human GSE30242 sample 2631 | 49/315 | 2.83E-17 | 4.23E-15 | 4.94 |
| 7 | metformin DB00331 human GSE33612 sample 2483 | 50/343 | 2.07E-16 | 2.65E-14 | 4.58 |
| 8 | etanercept DB00005 human GSE41663 sample 2605 | 45/322 | 3.29E-14 | 3.69E-12 | 4.33 |
| 9 | cisplatin DB00515 human GSE47856 sample 3145 | 40/267 | 8.93E-14 | 8.91E-12 | 4.68 |
| 10 | Lipopolysaccharide 11970143 human GSE5504 sample 3486 | 35/224 | 9.25E-13 | 8.30E-11 | 4.89 |

See Table 7. The first and fourth one, resveratrol, inhibits HCoV-229E and SARS-CoV-2 coronavirus replication in vitro [33]. The second, third, and fifth one, carboplatin, was proposed as a result of drug repositioning [21]. The seventh one, lipopolysaccharide, is listed in Table 6, too.

**Table 7.**
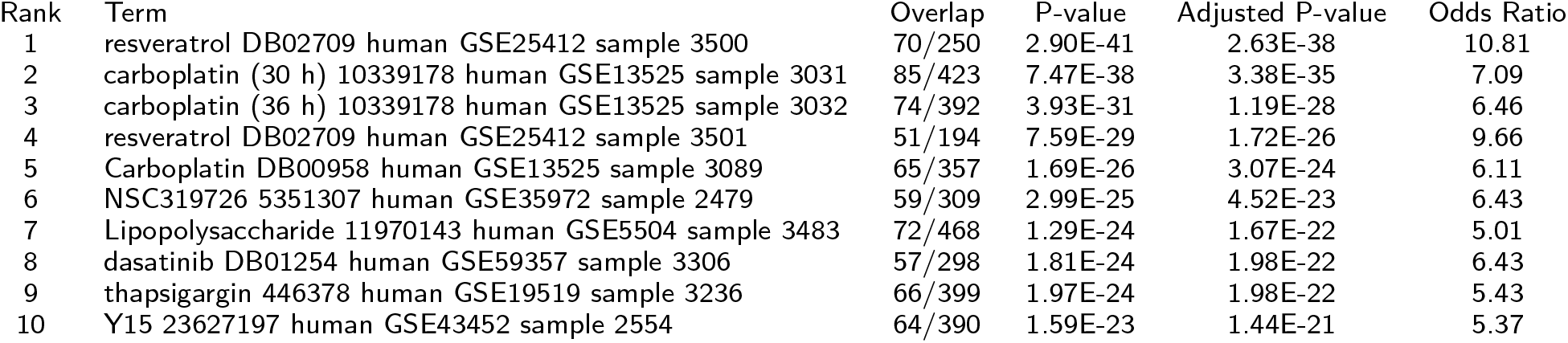
Drug perturbations from GEO up for A549 by DESeq2

The proposed method can predict effective drugs for COVID-19 based on gene expression analysis, at least, comparatively to DESeq2. Nevertheless, DESeq2 has less significance and has a tendency to list the same compounds multiple times. The proposed method can identify more convincing and diverse candidate compounds than DESeq2.

Based on the overlap between human genes known to interact with SARS-CoV-2 proteins and selected genes (Fig. 14) and from the point of drug repositioning, TD-based unsupervised FE with optimized SD is, at least, competitive with DESeq2.

### Comparison of methods using multi-organ measurements with multiple drug treatments

One might wonder if the proposed methods, TD- and PCA-based unsupervised FE with optimized SD, are applicable to a more complicated set-up. To investigate this point, we checked the case where multiple drugs are applied to mice whose gene expression of multiple tissues are measured, to which we applied TD-based unsupervised FE [34].

#### Enrichment of tissue-specific genes

In the previous study [34], although we applied TD-based unsupervised FE to gene expression profiles, there existed some problems. First of all, the number of genes selected was too small to have no false negatives. Using the optimized SD, the number of selected genes increased (Table 8; for more details, e.g., the definition of the four gene sets, neurons and testis, muscle, gastrointestine 1 and 2, see the previous study [34]. This topic has not been discussed herein as it is not directly related to the comparison of the performance between the original TD-based unsupervised FE and that with the optimised SD. The full list of the selected genes is available in Additional file 9). Although an increased number of genes is meaningless if the biological reliability is less, the biological reliability of selected genes is also improved (lower panel of Fig. 15, which corresponds to a present study and is associated with a greater number of cell lines and tissue specificity than that in the upper panel of Fig. 15, which corresponds to a previous study). Thus, the employment of optimized SD is also effective to a more complicated data set than simple pairwise comparisons between the treated and control samples investigated in the previous sections.

**Table 8.** Comparison of selected genes between TD-based unsupervised FE [34] and optimal SD with multi-organ data sets

| adjusted *P*-values | TD-based unsupervised FE [34] | TD-based unsupervised FE with optimized SD |  |
| --- | --- | --- | --- |
| | $\leq 0.01$ | $\leq 0.01$ | $\leq 0.1$ |
| Neuron | 18 | 356 | 472 |
| Muscle | 51 | 547 | 663 |
| Gastrointestine 1 | 97 | 1026 | 1322 |
| Gastrointestine 2 | 128 | 574 | 722 |

**Figure 15.**
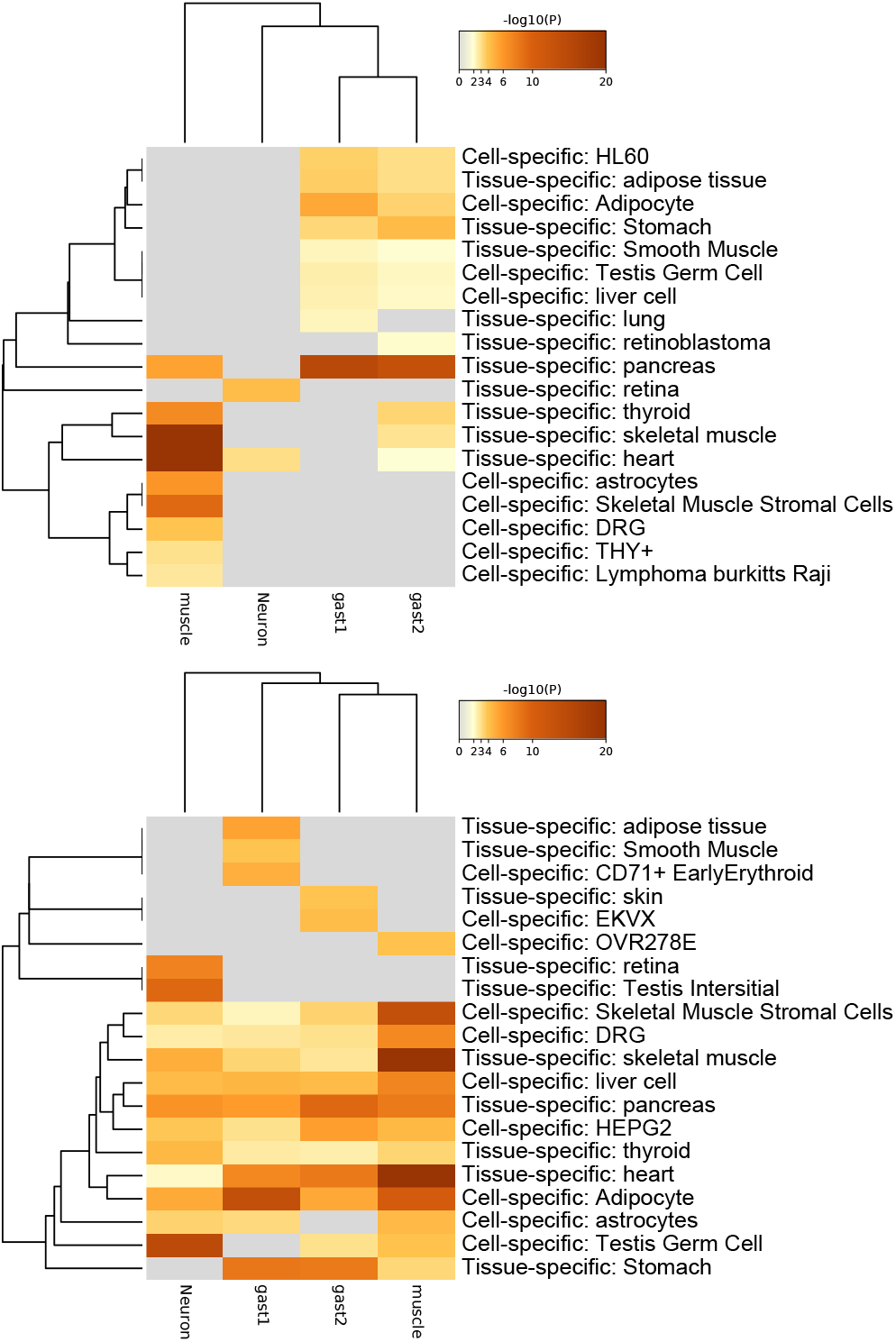
Enrichment analysis of cell and tissue specificity with Metascape [35]. Upper: original TD-based unsupervised FE (using genes with adjusted *P* ≤ 0.01 in Table **??**), lower; the present study with optimized SD (using genes with adjusted *P* ≤ 0.1 in Table 8).

#### Coincidence with drug treatment

We have also performed additional validation of the genes selected by TD-based unsupervised FE with optimized SD associated with adjusted *P*-values less than 0.1 (Table 8, full list is available in Additional files 10–13). We have uploaded selected genes to Enrichr [36] and evaluated the overlaps between the genes selected and those whose expression wasaltered with the treatment of the 15 drugs used in this study. Then, we found that all four gene sets in Table 8 had a significant overlap with the genes whose expression was altered with the treatment of 5 of the drugs (acetaminophen, cisplatin, clozapine, doxycycline, and olanzapine) in DrugMatrix, which does not include other drug treatments (Supplementary material). This suggests that TD-based unsupervised FE with optimal SD can correctly recognize drug treatments based on gene expression; this was impossible in the previous study [34] because of the very small number of genes selected (Table 8). Thus, considering the optimization of SD enables TD-based unsupervised FE to recognize a greater number of biologically reliable genes than the original TD-based unsupervised FE, which did not include the optimization of SD.

## Discussion

In this study, we have introduced the optimization of SD to TD- and PCA-based unsupervised FE and have improved their performance by increasing the identified DEGs associated with greater biological reliability. One of the striking features is that DEGs with lesser gene expression are less likely recognized even with the same LFC, if the genes are selected by TD- and PCA-based unsupervised FE with optimized SD. In DESeq2, the tendency that less expressed genes are hardly recognized as DEGs is artificially introduced by assuming dispersion relation, eq. (1). Nevertheless, in PCA- and TD-based unsupervised FE, it is automatically introduced. Generally, there exists a relationship between difference, Δ of two variables, *x* and *y*, and LFC as

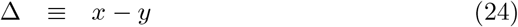

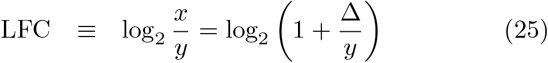

Then

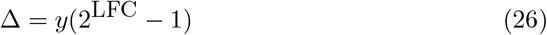

Because *v*_2*j*_ (Fig. 2B) corresponds to Δ, if DEGs are identified using *u*_2*i*_ that corresponds to *v*_2*j*_ as in TD- and PCA-based unsupervised FE (see eqs. (6) and (12)), DEGs associated with the same LFC are less likely selected for the smaller *y* that corresponds to *µ*. This results in the distribution of DEGs in MAPlot (Fig. 8), where genes with the same LFC (vertical axis) are less likely identified as DEGs with smaller gene expression (horizontal axis). Figure 16 shows the MAPlot drawn using two independent random variables obeying the same positive uniform distribution; the red colored region associated with |Δ| larger than some threshold values qualitatively represents the tendency that indicates that a smaller *x* + *y* is less likely selected even with the same LFC, 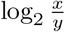. Thus, TD- and PCA-based unsupervised FE can introduce the tendency that genes with less expression are less likely to be DEGs, even with the same amount of LFC more naturally than DESeq2, which has to manually introduce a dispersion relation, eq. (1).

**Figure 16.**
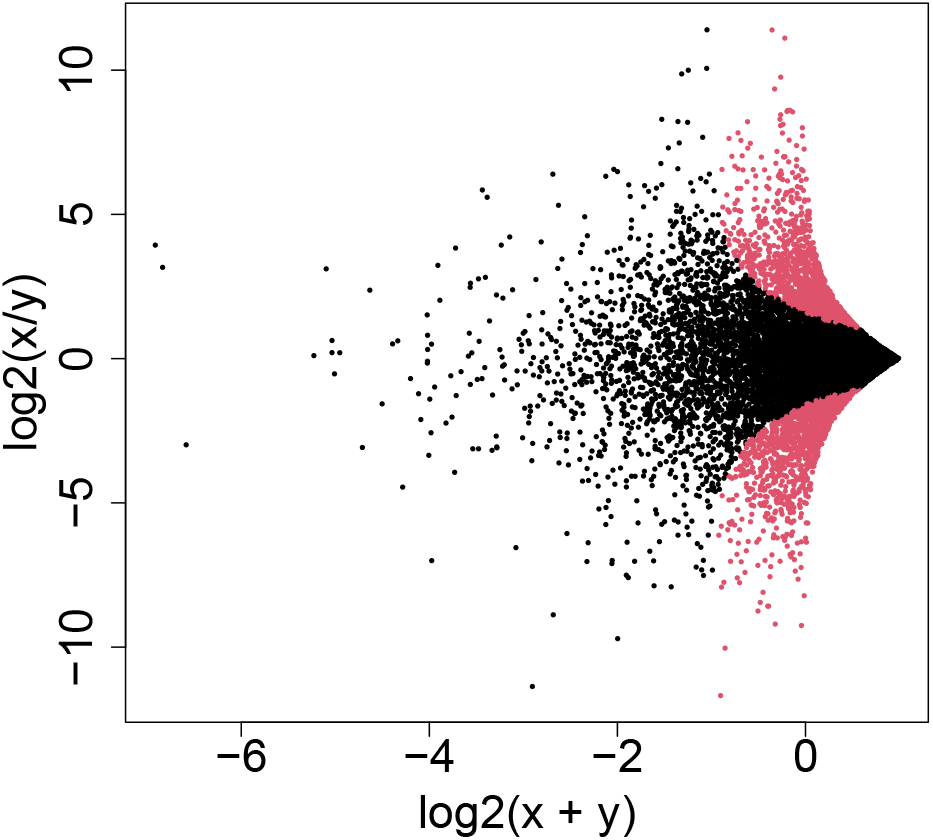
“MAPlot” using two independent variables, *x* and *y*, drawn from uniform distribution ∈ [0, 1]. Red dots are associated with |*x* − *y*| *>* 0.5.

In addition to this, although DESeq2 assumes NB distribution that does not have any rationalization other than that it takes only positive values and has a tunable mean as well as variance simultaneously, TD- and PCA-based unsupervised FE assume only that *u*_*ℓi*_ obeys the Gaussian distribution (eqs. (6) and (12)), which is more reasonable because Gaussian distributions can generally appear when independent random variables are summed up. Actually, NOISeq does not assume NB distribution as well but achieves comparative performance with DESeq2 (Fig. 13). In this sense, TD- and PCA-based unsupervised FE can realize DEG distribution in an MAPlot more naturally than DE-Seq2.

Another remarkable point of TD- and PCA-based unsupervised FE with optimized SD is that it does not have to screen for selected genes by LFC after the genes are selected using *P*-values. As can be seen in Fig. 10, state-of-art methods, including DESeq2, often identify too many DEGs. In these circumstances, LFC is often used to reduce the number of DEGs. Nevertheless, Stupnikov et al [37] found that the coincidence of the selected genes among the various state-of-art methods drastically decreases if the genes selected based on *P*-values are further screened with LFC. In this sense, TD- and PCA-based unsupervised FE with optimized SD are more promising methods than state-of-art methods that need screening by LFC to yield a reasonable number of DEGs.

Yet another advantage is that TD- and PCA-based unsupervised FE have already been applied to a wide range of problems. Not only can optimized SD improve the performance of PCA- and TD-based unsupervised FE, as can be seen in Figs. 14 and 15, but also the alteration is limited to the last stage, i.e., *P*-value computation, eqs. (6) and (12). Thus, the optimized SD is expected to improve the performance in a wide range of problems, to which TD- and PCA-based unsupervised FE have been applied.

## Conclusions

In this study, we optimized SD to improve TD- and PCA-based unsupervised FE. As a result, not only the obtained DEGs increased and became reasonable in number but also the histogram of 1-*P* became more reliable, i.e., more coincident with the null hypothesis that SVV and PC obey Gaussian distribution. In addition to this, TD- and PCA-based unsupervised FE provide reliable distribution of DEGs in MAPlot, i.e., less expressed genes are less likely selected as DEGs even if they are associated with the same LFC; this property was implemented manually by assuming dispersion relation, eq.(1), in DESeq2. The biological reliability of the selected genes is also much better by this method than by other state-of-art methods. These points suggest that TD- and PCA-based unsupervised FE are superior than state-of-art methods in terms of achieving better performance with less assumption.

## Methods

### Gene expression profiles

#### MAQC

Seven human brain expression profiles were downloaded from SRA [38] (ID SRX016359), and seven UHR expression profiles were downloaded from SRA (ID SRX016367). Fourteen FASTQ files were mapped to the hg38 human genome using rapmap [39]. htseqcount [40] was used to convert the obtained bam files to count data files using the gtf file taken from ftp://ftp.ensembl.org/pub/release-105/gtf/homo_sapiens/Homosapiens.GRCh38.105.gtf.gz

#### SEQC

SEQC [13] were obtained from bioconductor [41] as an experimental package, seqc. It includes thirteen profiles shown in Fig. 11. For more details, see Vignettes in the seqc experimental package.

### The histogram composed of Gaussian distribution and outliers in Fig. 4

The Gaussian part is one thousand values drawn from Gaussian distribution with zero mean and an SD of one. Outliers are 100 values, which are equal to 5.

### PCA-based unsupervised FE applied

#### MAQC

Genes not expressed in any of the 14 samples have been excluded. Four rows having annotations “no feature”, “ambiguous”, “not aligned”, and “alignment not unique” have also been excluded. As a result, we got *x*_*ij*_ ∈ ℝ^40933×14^. The *x*_*ij*_ was processed as described in the main text.

#### SEQC

Regardless of which of the 13 data sets was considered, only those genes expressed in all samples were considered. An individual data set has a distinct number of rows (genes) and columns (samples). The *x*_*ij*_ obtained from an individual data set was processed as described in the main text.

#### SARS-CoV-2

All processes used were exactly the same as those described in the previous study [14]. After obtaining *u*_5*i*_, the SD was optimized as described in the main text.

#### Multi-organ

All processes used were exactly the same as those described in the previous study [34]. After getting *u*_*ℓi*_, the SD was optimized as described in the main text.

### Optimization of SD

At first, a histogram of 1− *P*_*i*_ was computed using hclust function in R with the “break=100” option. Then, an SD of the binned histogram, hc$count associated with hc$breaks less than 1-*P* whose adjusted *P*-value was less than threshold value *P*_0_, was minimized using optim function in R. The R code has been provided in additional file 14 to show how to optimize SD in an individual data set.

### Coincidence between PCA-based unsupervised FE and DESeq2

The coincidence between PCA-based unsupervised FE HLDESeq2 was evaluated by AUC (Fig. 9) as follows.

At first, the top 1000 genes based on *P*-values computed by DESeq2 were regarded positive and the remaining genes were regarded negative. Then, *P*-values computed by PCA-based unsupervised FE were used to predict positive genes. Using this result, AUC was computed. Next, on the contrary, the top 1000 genes based on *P*-values computed by PCA-based unsupervised FE were regarded positive and the remaining genes were negatives. Then, *P*-values computed by DESeq2 were used to predict positive genes. Using this result, AUC was computed.

### Enrichment analyses

Enrichment analyses were performed using either Metascape [35] or Enrichr [36] by uploading gene symbols. If the gene ID was not a gene symbol in individual data sets, the gene ID conversion tool in Database for Annotation, Visualization, and Integrated Discovery (DAVID) [42, 43] was used for conversion.

### DEG identification of SARS-CoV-2 data by DESeq2

We used author-provided adjusted *P*-values and LFC (in supplementary data in their paper) to identify DEGs. If we considered only adjusted *P*-values to identify DEGs, DESeq2 would identify too many genes (Table 9). Thus, we had to consider LFC as well. Table 9 shows the number of DEGs used in this study. The evaluation of the overlap with human genes known to interact with SARS-CoV-2 proteins is available in Supplementary materials. The best one, that for the ACE2-expressed A549 cell line, is also included in the main text as Fig. 14.

**Table 9.**
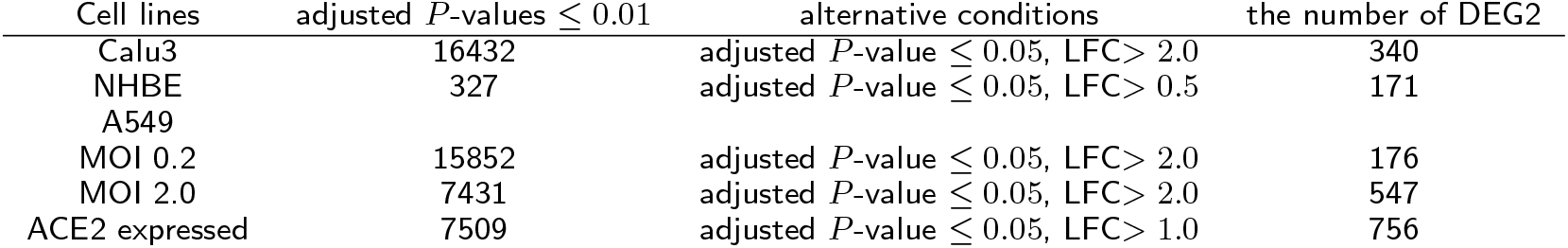
The number of DEGs in SARS-CoV-2 study by DESeq2 (based on author-provided supplementary material)

## Declarations

Ethics approval and consent to participate

Not applicable.

Consent for publication

Not applicable.

## Availability of data and materials

The MAQC data set can be downloaded from SRA with ID SRX016359 and SRX016367. The SEQC data is a part of the bioconductor seqc package. SARS-CoV-2 data can be downloaded from Gene Expression Omnibus (GEO) with the GEO ID GSE147507. Multi-organ data can be downloaded from GEO with the GEO ID GSE142068.

## Author’s contributions

YHT planned the research and performed analyses. YHT and TT evaluated the results, discussions, and outcomes and drafted and reviewed the manuscript.

## Acknowledgements

Not applicable.

## Additional Files

Additional file 1 — Genes selected by DESeq2 for MAQC

Genes associated with adjusted *P*-values less than 0.1 using DESeq2 and enrichment analysis associated with them.

Additional file 2 — Genes selected by PCA-based unsupervised FE with optimized SD for MAQC

Genes associated with adjusted *P*-values less than 0.1 using PCA-based unsupervised FE with optimized SD and enrichment analysis associated with them.

Additional file 3 — Genes selected by EdgeR for MAQC

Genes associated with adjusted *P*-values less than 0.1 using EdgeR and enrichment analysis associated with them.

Additional file 4 — Genes selected by voom for MAQC

Genes associated with adjusted *P*-values less than 0.1 using voom and enrichment analysis associated with them.

Additional file 5 — Genes selected by NOISeq for MAQC

Genes associated with adjusted *P*-values less than 0.1 using NOISeq and enrichment analysis associated with them.

Additional file 6 — Genes selected by TD-based unsupervised FE with optimized SD for SARS-CoV-2

Genes associated with adjusted *P*-values less than 0.1 by TD-based unsupervised FE with optimized SD for SARS-CoV-2 and drug repositioning associated with the genes.

Additional file 7 — Overlap with human genes known to interact with SARS-CoV-2 protein by DESeq2

Overlap with human genes known to interact with SARS-CoV-2 proteins by DESeq2.

Additional file 8 — Genes selected for SARS-CoV-2 infected A549 cell lines by DESeq2

Genes selected by DESeq2, for A549 cell lines, shown in Table 14, and drug repositioning.

Additional file 9 — Genes selected by TD-based unsupervised FE with optimized SD for multi-organ study

Genes selected by TD-based unsupervised FE with optimized SD for a multi-organ study.

Additional file 10 — Drug repositioning for neuron and tesis gene sets Drug repositioning for neuron and tesis gene sets.

Additional file 11 — Drug repositioning for muscle gene sets Drug repositioning for muscle gene sets.

Additional file 12 — Drug repositioning for gast 1 gene sets Drug repositioning for gast 1 gene sets.

Additional file 13 — Drug repositioning for gast 2 gene sets Drug repositioning for gast 2 gene sets.

Additional file 14 — Source code

R source code to perform PCA- and TD-based unsupervised FE with optimized SD.

