## Supplementary material for "Tensor decomposition- and principal component analysis-based unsupervised feature extraction to select more reasonable differentially expressed genes: Optimization of standard deviation versus state-of-art methods": Additional_file_14.R

```
#MAQC
#mapping reads to hg38 (ourside R)
rapmap quasiindex -t genome.fa -i ref_index >
rapmap quasimap -i ~/RapMap/bin/ref_index -r <(gunzip -c SRR037477.fastq.gz) -t 8 -o SRR037477_mapped_reads.sam > nohup_rapmap_SRR037477.out
samtools view -bS SRR037477_mapped_reads.sam > SRR037477_mapped_reads.bam
samtools sort SRR037477_mapped_reads.bam > SRR037477_mapped_reads_sort.bam 
samtools index SRR037477_mapped_reads_sort.bam  
#convert count 
htseq-count -f bam SRR037440_mapped_reads_sort_cottected.bam Homo_sapiens.GRCh38.105.gtf > SRR037440.txt 
#inside R
#suppose count data is in current.directoyy
Files = list.files("./",pattern=".txt")
x_all <- NULL
for (i in c(1:length(Files)))
{
    cat(i, " ")
    x <- read.csv(Files[i],sep="\t",header=F)
    x_all <- cbind(x_all,x[,2])
}
x_all<- data.frame(x[,1],x_all)
x_all <-x_all[rowSums(x_all[,-1])!=0,]
x_all <- x_all[1:40933,]
x_all[,-1]<- scale(x_all[,-1])
PCA <- prcomp(scale(x_all[,-1]))

th <- function(sd){
    P2 <- pchisq((PCA$x[,2]/sd)^2,1,lower.tail=F)
    hc<- hist(1-P2,breaks=100,plot=F)
    return(sd(hc$count[1:sum(hc$breaks<1-min(P2[p.adjust(P2,"BH")>0.1]))]))
}

sd <- optim(0.8,th,method="Brent",lower=0,upper=1)$par
P2 <- pchisq((PCA$x[,2]/sd)^2,1,lower.tail=F)
genes <- x_all[p.adjust(P2,"BH")<0.1,1]

#SEQC
require(seqc)
#remove coment one of them
#LAB <- ILM_refseq_gene_BGI #ERCC 43, 14261 genes, 80 vs 80 samples 
#LAB <- ILM_refseq_gene_AGR #ERCC 51, 15601 genes, 64 vs 64 samples
#LAB <- ILM_refseq_gene_CNL #ERCC 42, 14028 genes, 75 vs 75 samples 
#LAB <- ILM_refseq_gene_COH #ERCC 48, 16224 genes, 32 vs 32 samples
#LAB <- ILM_refseq_gene_MAY #ERCC 43, 13753 genes, 80 vs 80 samples
#LAB <- ILM_refseq_gene_NVS #ERCC 51, 15600 genes, 64 vs 64 samples
#LAB <- LIF_refseq_gene_LIV #ERCC 61, 16135 genes, 10 vs 10 samples
#LAB <- LIF_refseq_gene_NWU #ERCC 43, 11842 genes, 60 vs 60 samples
#LAB <- LIF_refseq_gene_PSU #ERCC 40, 10745 genes, 60 vs 60 samples
#LAB <- LIF_refseq_gene_SQW #ERCC 44, 11659 genes, 60 vs 60 samples
#LAB <- ROC_refseq_gene_MGP #ERCC 32, 11055 genes, 2 vs 2 samples 
#LAB <- ROC_refseq_gene_NYU #ERCC 36, 11672 genes, 2 vs 2 samples
#LAB <- ROC_refseq_gene_SQW #ERCC 34, 10613 genes, 2 vs 2 samples
index_A <- substring(colnames(LAB),1,1)=="A"
index_B <- substring(colnames(LAB),1,1)=="B"
x_all <- data.frame(ILM_refseq_gene_BGI[,1:4],cbind(LAB[,index_A],LAB[,index_B]))

x_all <- x_all[rowSums(x_all[,-c(1:4)]!=0)==dim(x_all[,-c(1:4)])[2],] #remove zeros

PCA <- prcomp(scale(x_all[,-c(1:4)]))

th <- function(sd){
    P2 <- pchisq((PCA$x[,2]/sd)^2,1,lower.tail=F)
    hc<- hist(1-P2,breaks=100,plot=F)
    return(sd(hc$count[1:sum(hc$breaks<1-min(P2[p.adjust(P2,"BH")>0.1]))]))
}
#remove comment one of them
#sd <- optim(0.5,th,method="Brent",lower=0,upper=1)$par #other than MGR,ROC_SQW
#sd <- optim(0.2,th)$par #for MGR
#sd <- optim(0.1,th)$par #for ROC_SQW
P2 <- pchisq((PCA$x[,2]/sd)^2,1,lower.tail=F)
genes <- x_all[p.adjust(P2,"BH")<0.1,1]

#SARS-CoV-2
#dowlaod file from 
#https://ftp.ncbi.nlm.nih.gov/geo/series/GSE147nnn/GSE147507/suppl/GSE147507_RawReadCounts_Human.tsv.gz
x <- read.csv("GSE147507_RawReadCounts_Human.tsv.gz",sep="\t",comment.char="!")

Z <- array(NA,c(dim(x)[1],5,2,3))
Z[,1,1,] <- data.matrix(x[,-1][,33:35]) #Calu3
Z[,1,2,] <- data.matrix(x[,-1][,36:38]) #Calu3
Z[,2,1,] <- data.matrix(x[,-1][,1:3]) #NHBE
Z[,2,2,] <- data.matrix(x[,-1][,4:6]) #NHBE
Z[,3,1,] <- data.matrix(x[,-1][,7:9]) #A549 MOI3
Z[,3,2,] <- data.matrix(x[,-1][,10:12]) #A540 MOI3 
Z[,4,1,] <- data.matrix(x[,-1][,21:23])
Z[,4,2,] <- data.matrix(x[,-1][,24:26])
Z[,5,1,] <- data.matrix(x[,-1][,27:29])
Z[,5,2,] <- data.matrix(x[,-1][,30:32])
Z <- apply(Z,2:4,scale)
require(rTensor)
HOSVD <- hosvd(as.tensor(Z),c(10,5,2,3))
    
th <- function(sd){
    P2 <- pchisq((HOSVD$U[[1]][,5]/sd)^2,1,lower.tail=F)
    hc<- hist(1-P2,breaks=100,plot=F)
    return(sd(hc$count[1:sum(hc$breaks<1-min(P2[p.adjust(P2,"BH")>0.1]))]))
}
sd <- optim(0.001,th)$par
P2 <- pchisq((HOSVD$U[[1]][,5]/sd)^2,1,lower.tail=F)

genes <- x[p.adjust(P2,"BH")<0.1,1]

#multi organ
#download files from
#https://www.ncbi.nlm.nih.gov/geo/query/acc.cgi?acc=GSE142068
#GSE142068_count_AdrenalG.txt.gz  GSE142068_count_PituitaryG.txt.gz
#GSE142068_count_Aorta.txt.gz     GSE142068_count_SkMuscle.txt.gz
#GSE142068_count_BM.txt.gz        GSE142068_count_Skin.txt.gz
#GSE142068_count_Brain.txt.gz     GSE142068_count_Skull.txt.gz
#GSE142068_count_Colon.txt.gz     GSE142068_count_Spleen.txt.gz
#GSE142068_count_Eye.txt.gz       GSE142068_count_Stomach.txt.gz
#GSE142068_count_Heart.txt.gz     GSE142068_count_Testis.txt.gz
#GSE142068_count_Ileum.txt.gz     GSE142068_count_Thymus.txt.gz
#GSE142068_count_Jejunum.txt.gz   GSE142068_count_ThyroidG.txt.gz
#GSE142068_count_Kidney.txt.gz    GSE142068_count_WAT.txt.gz
#GSE142068_count_Liver.txt.gz     GSE142068_count_Lung.txt.gz      
#GSE142068_count_Pancreas.txt.gz  GSE142068_count_ParotidG.txt.gz 

files <- list.files("./",pattern="txt.gz")
names<-NULL
for (i in c(1:length(files)))
{
    cat(i," ")
    x <- read.csv(files[i],sep="\t",header=T)
    TABLE<- table(gsub(".rep4","",gsub(".rep3","",gsub(".rep2","",gsub(".rep1","",colnames(x))))))
    names <- unique(c(names,names(TABLE)))
}
TABLES <- matrix(0,length(names),1)
TABLES<-data.frame(names,TABLES)
colnames(TABLES) <-c("Var1","Freq")
for (i in c(1:length(files)))
{
    cat(i," ")
    x <- read.csv(files[i],sep="\t",header=T)
    TABLE<- table(gsub(".rep4","",gsub(".rep3","",gsub(".rep2","",gsub(".rep1","",colnames(x))))))
    TABLES <- merge(TABLES,data.frame(TABLE),by="Var1")
}
TABLES <- TABLES[-10,-2]

Z <- array(NA,(c(dim(x)[1],24,18,2)))
for (i in c(1:length(files)))
{
    cat(i," ")
    x <- read.csv(files[i],sep="\t",header=T)
    sample_name <- gsub(".rep4","",gsub(".rep3","",gsub(".rep2","",gsub(".rep1","",colnames(x)))))
    index <- outer(sample_name,TABLES[,1],"==")
    for( j in c(1:18))
    {
        Z[,i,j,] <- data.matrix(x[,index[,j]][,1:2])
    }
}
Z <- apply(Z,c(2:4),scale)
require(rTensor)
HOSVD <- hosvd(as.tensor(Z),c(10,24,18,2))

#remove comment one of the comments
#k<-2 #Neuron
#k<-4 #Muscle
#k<-5 #Gast 1
#k<-6 #Gast2
th <- function(sd){
    P2 <- pchisq((HOSVD$U[[1]][,k]/sd)^2,1,lower.tail=F)
    hc<- hist(1-P2,breaks=100,plot=F)
    return(sd(hc$count[1:sum(hc$breaks<1-min(P2[p.adjust(P2,"BH")>0.1]))]))
}
sd <- optim(0.001,th)$par
P2 <- pchisq((HOSVD$U[[1]][,k]/sd)^2,1,lower.tail=F)

genes <- x[p.adjust(P2,"BH")<0.1,1]
    
```
